# Insights on the regulation and function of the CRISPR/Cas transposition system located in the pathogenicity island VpaI-7 from *Vibrio parahaemolyticus* RIMD2210633

**DOI:** 10.1101/2025.03.26.645530

**Authors:** Jesús E. Alejandre-Sixtos, Kebia Aguirre-Martínez, Jessica Cruz-López, Aliandi Mares-Rivera, Samanda Álvarez-Martínez, David Zamorano-Sánchez

## Abstract

CRISPR/Cas mediated transposition is a recently recognized strategy for horizontal gene transfer in a variety of bacterial species. However, our understanding of the factors that control their function in their natural hosts is still limited. In this work we report our initial genetic characterization of the elements associated with the CRISPR/Cas-transposition machinery (CASTm) from *Vibrio parahaemolyticus* (*Vpa*CASTm), which are encoded within the pathogenicity island VpaI-7. Our results revealed that the components of the *Vpa*CASTm and their associated CRISPR arrays (*Vpa*CAST system) are transcriptionally active in their native genetic context. Furthermore, we were able to detect the presence of polycistrons and several internal promoters within the loci that compose the *Vpa*CAST system. Our results also suggest that the activity of the promoter of the atypical CRISPR array is not repressed by the baseline activity of its known regulator VPA1391 in *V. parahaemolyticus*. Additionally, we found that the activity of the promoter of *tniQ* was modulated by a regulatory cascade involving ToxR, LeuO and H-NS. Since it was previously reported that the activity of the *Vpa*CAST system was less efficient than that of the *Vch*CAST system at promoting transposition of a miniaturized CRISPR-associated transposon (mini-CAST) in *Escherichia coli*, we analyzed if the transposition efficiency mediated by the *Vpa*CAST system could be enhanced inside its natural host *V. parahaemolyticus*. We provide evidence that this might be the case suggesting that there could be host induction factors in *V. parahaemolyticus* that could enable more efficient transposition of CASTs.

**Importance:** Mobile genetic elements such as transposons play important roles on the evolutionary trajectories of bacterial genomes. The success of transposon dissemination depends on their ability to carry selectable markers that improve the fitness of the host cell or loci with addictive traits such as the toxin-antitoxin systems. Here we aimed to characterize a transposon from *Vibrio parahaemolyticus* that carries and could disseminate multiple virulence factors. This transposon belongs to a recently discovered family of transposons whose transposition is guided by crRNA. We showed that the transposition machinery of this transposon is transcribed in *V. parahaemolyticus* and that there are likely host associated factors that favor transposition in the natural host *V. parahaemolyticus* over transposition in *Escherichia coli*.

## Introduction

*Vibrio parahaemolyticus* is a cosmopolitan bacterium capable of colonizing the intestine of marine animals such as shrimp and causing gastroenteritis in humans (1). Its virulent properties are mainly associated with the production of three hemolysins (TDH, TRH and TLH) and two type 3 secretion systems, one located on chromosome 1 (T3SS1) and another on chromosome 2 (T3SS2) (2, 3). Two copies of the thermostable direct hemolysin and the T3SS2 are encoded in the pathogenicity island VpaI-7 (2) (Fig. 1A). It has been proposed that this island was acquired by horizontal gene transfer (HGT) recently in the evolutionary history of the pandemic clone of *V. parahaemolyticus* O3:K6 during a pandemic outbreak that occurred between the years 1995 and 1996 (4–6). The continued dissemination of virulence factors through the horizontal transfer of VpaI-7 could result in the emergence of bacterial strains with greater pathogenic capacity (7, 8). To date, the molecular mechanisms that allowed the spread of the VpaI-7 island in *V. parahaemolyticus* strains remain unknown as well as whether it retains the ability to move by horizontal transfer.

**Figure 1.**
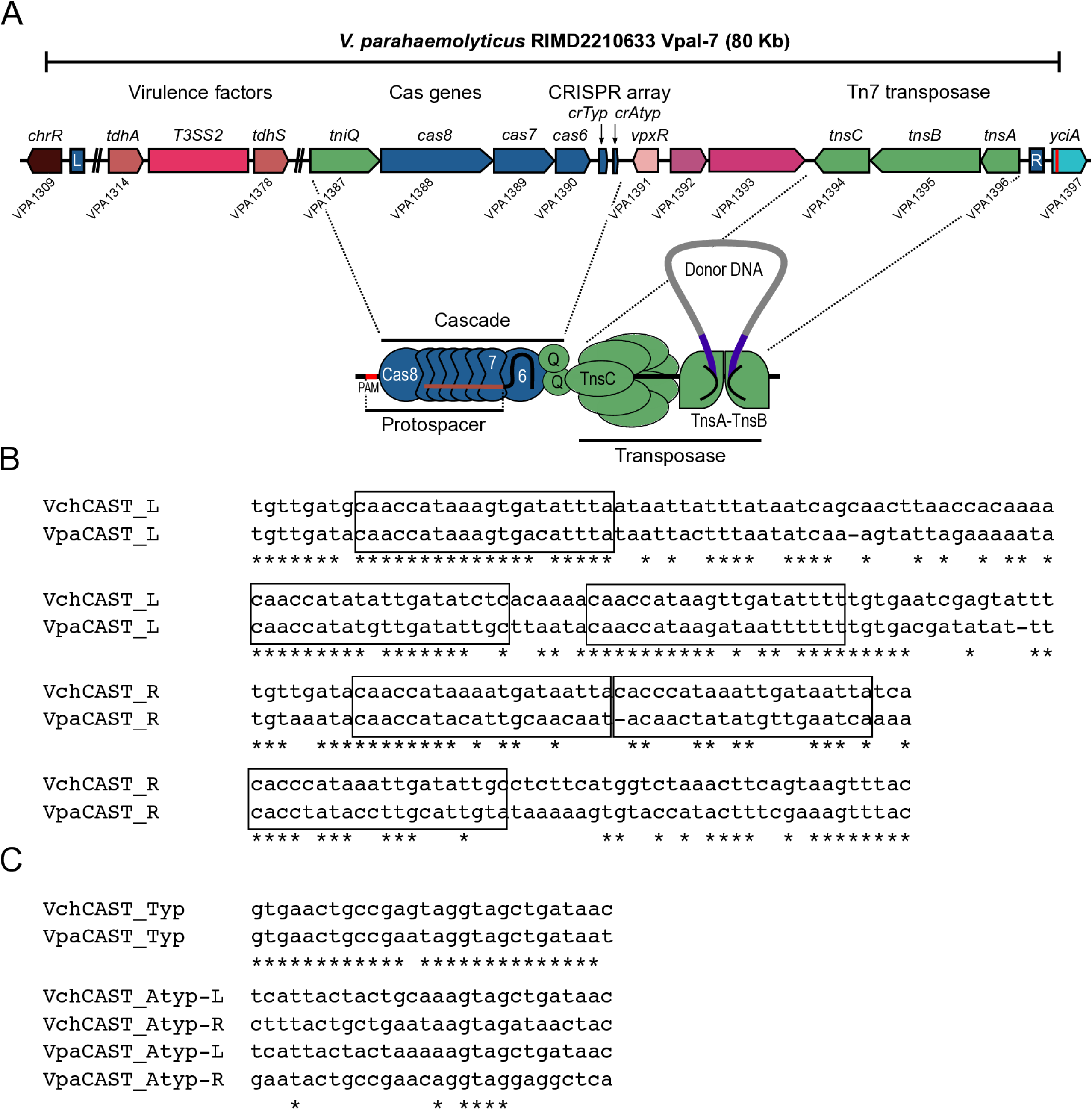
The pathogenicity island VpaI-7 from *V. parahaemolyticus* RIMD2210633 is within a CRISPR/Cas-transposon. A) Diagram depicting genes encoding relevant components of the VpaI-7 island including virulence factors and the elements of a Type IF3 CRISPR/Cas transposition system. B) Sequence alignment of the predicted TnsB binding sites (ToxR-box) in the L and R-terminal regions of the *Vch*CAST and *Vpa*CAST systems. The ToxR-boxes are delimited by rectangles. C) Sequence alignment of the direct repeats flanking the spacer sequences of the typical and atypical CRISPR arrays from the *Vch*CAST and *Vpa*CAST systems.

Recently, a series of transposons of the Tn7 family whose transposition mechanism involves the joint activity of a TnsABC transposase, a TniQ family protein and a minimalistic CRISPR/Cas system, have been characterized (9). These types of transposons are commonly referred to as “CRISPR-associated transposons” (CASTs) (10–12) and are widely distributed among a vast variety of bacterial genera including the *Vibrio* genus (13, 14). Minimalistic CRISPR/Cas systems, unlike canonical CRISPR/Cas systems, lack the proteins necessary for CRISPR array adaptation and target interference, processes involved in immune memory mechanisms and defense against foreign DNA, respectively (13). The minimalist CRISPR/Cas systems encoded within CASTs have the function of guiding the transposition of these mobile genetic elements through the base complementarity of a CRISPR RNA (crRNA) with their double strand DNA target site known as the protospacer sequence (15) (Fig. 1A). The transposition of CASTs encoding minimalistic CRISPR/Cas I-F systems depends on the CRISPR-associated complex for antiviral defense (Cascade) composed of the Cas8, Cas7 and Cas6 proteins (16) (Fig. 1A). The Cascade I-F recognizes a 5’-CC protospacer-adjacent motif (PAM), and the crRNA forms a DNA-RNA double strand through base complementarity with the protospacer to promote the transposon insertion event (17) (Fig. 1A). Once the Cascade complex recognizes its target, it uses TniQ as an intermediary to recruit the transposase composed of TnsA, TnsB and TnsC (16) (Fig. 1A). The TnsA-TnsB complex recognizes the asymmetric right (R) and left (L)-terminal sequences, which have multiple TnsB binding sites and delimit the CAST, while TnsC acts as a regulator of transposition by acting as a bridge between TniQ and TnsA-TnsB (18–22) (Fig. 1).

Sequence analysis of the VpaI-7 island strongly suggests that it is a CAST, it is flanked by Tn7like R and L-terminal sequences and encodes the proteins TnsA (VPA1396), TnsB (VPA1395) TnsC (VPA1394), TniQ (VPA1387) and a minimalistic CRISPR/Cas system I-F3a composed of the proteins Cas8 (VPA1388), Cas7 (VPA1389) and Cas6 (VPA1390), as well as two independent CRISPR arrays, named “typical” and “atypical” (23) (Fig. 1). The R and Lterminal sequences have putative TnsB-boxes that share sequence identity with the TnsBboxes previously characterized for the *Vch*CAST system encoded in the Tn6677 transposon from *V. cholerae* HE-45 and homologous Type I-F3 CAST systems (24). The typical CRISPR array of VpaI-7 is composed of three direct repeat sequences and two spacer sequences complementary to plasmid sequences found in various species of the *Vibrio* genus (25, 26).

The atypical CRISPR array contains two imperfectly repeated sequences flanking a single spacer (23, 24). The spacer of the atypical array has sequence complementarity with the *yciA* gene (VPA1397) (23, 24), which is the gene located downstream of the R-terminal sequence of the CAST on chromosome 2 from *V. parahaemolyticus* (Fig. 1A). It has been documented that CASTs typically transposed in an R-L orientation (16, 27). This suggests that the spacer of the atypical array allowed the insertion of the CAST harboring the VpaI-7 island.

The CAST transposition mechanism has been mainly studied in the heterologous background of *Escherichia coli* (16, 23, 24, 27), outside of its native context. The first CRISPRmediated transposition system studied experimentally was the one encoded in the Tn6677 transposon from *V. cholerae* HE-45 (16). The CRISPR/Cas-transposition machinery (CASTm) of this CAST was found to be functional in *E. coli* and was used to generate a genetic tool, named INTEGRATE, for the generation of programmed mini-CAST insertional mutations (27). The INTEGRATE tool is practical for studying CAST transposition since it consists of a single plasmid containing a programmable CRISPR array, the *tniQ*, *cas8*, *cas7*, *cas6*, *tnsA*, *tnsB* and *tnsC* genes, all under the control of the strong synthetic promoter P_J23119_ (27). An interchangeable mini-CAST genetic cargo flanked by the R and L-terminal sequences is present on the same plasmid (27). Using INTEGRATE-inspired approaches, previous studies reported that the CRISPR-associated transposition system encoded in VpaI-7 (*Vpa*CAST system) is functional in *E. coli*, although with lower efficiency than the system encoded in Tn6677 (23, 24, 27).

Little is known about the function and regulation of CAST systems such as the *Vpa*CAST system in their native genetic background. In this regard, it was recently reported that the product of the gene VPA1391, located downstream of the CRISPR arrays within VpaI-7, is a conserved Xre-type transcriptional regulator that exerts a negative effect on the expression of the atypical array (23). The evidence presented was obtained through biochemical experiments and genetic studies in the heterologous host *E. coli* (23), therefore the exact contribution of this regulator in controlling the expression of the atypical array crRNA (crAtyp) in its native genetic background is unknown so far. Furthermore, to our knowledge, there are no studies directly evaluating the transcriptional activity, regulation and arrangement of the CAST system elements of the VpaI-7 island.

In this work we asked some fundamental questions regarding the physiological relevance of the *Vpa*CAST system in *V. parahaemolyticus*. We asked 1) if the elements of the *Vpa*CAST system are transcribed under standard laboratory growth conditions; 2) if the active promoters of the components of the *Vpa*CAST system are subject of transcriptional regulation; and 3) if the components of the *Vpa*CASTm are equally efficient to promote transposition in its native genetic background as in the heterologous host *E. coli*. The observations and discoveries reported in this work will open new areas of inquiry that will lead to a better understanding of the contribution of CRISPR/Cas-mediated transposition to the evolution of *Vibrio* spp. pathogenicity and colonization success.

## Results

### The genes involved in CRISPR/Cas-mediated transposition are transcriptionally active in *V. parahaemolyticus* RIMD2210633

To begin to address the physiological relevance of the *Vpa*CAST system in *V. parahaemolyticus* RIMD2210633 we first evaluated if its components were transcriptionally active under standard laboratory growth conditions. The apparent organization of the genetic context of these genes suggests that *tniQ*, *cas8*, *cas7* and *cas6* are part of a 4 gene operon encoded on the leading strand, while *tnsA*, *tnsB* and *tnsC* are forming an operon encoded on the lagging strand (Fig. 2A). An RT-PCR analysis revealed the presence of transcripts corresponding to *tniQ*, *cas8*, *cas7*, *cas6*, *tnsA*, *tnsB* and *tnsC* (Fig. 2B). We also found potential polycistronic transcripts containing *tniQ*-*cas8*, *cas8*-*cas7*, *cas7*-*cas6*, *tnsA*-*tnsB*, and *tnsB*-*tnsC* (Fig. 2C). The amplification of the putative *tniQ*-*cas8*, *cas8*-*cas7* and *cas7-6* polycistronic units was weaker compared to the amplification of the polycistronic units corresponding to *tnsA*-*tnsB* and *tnsB*-*tnsC*.

**Figure 2.**
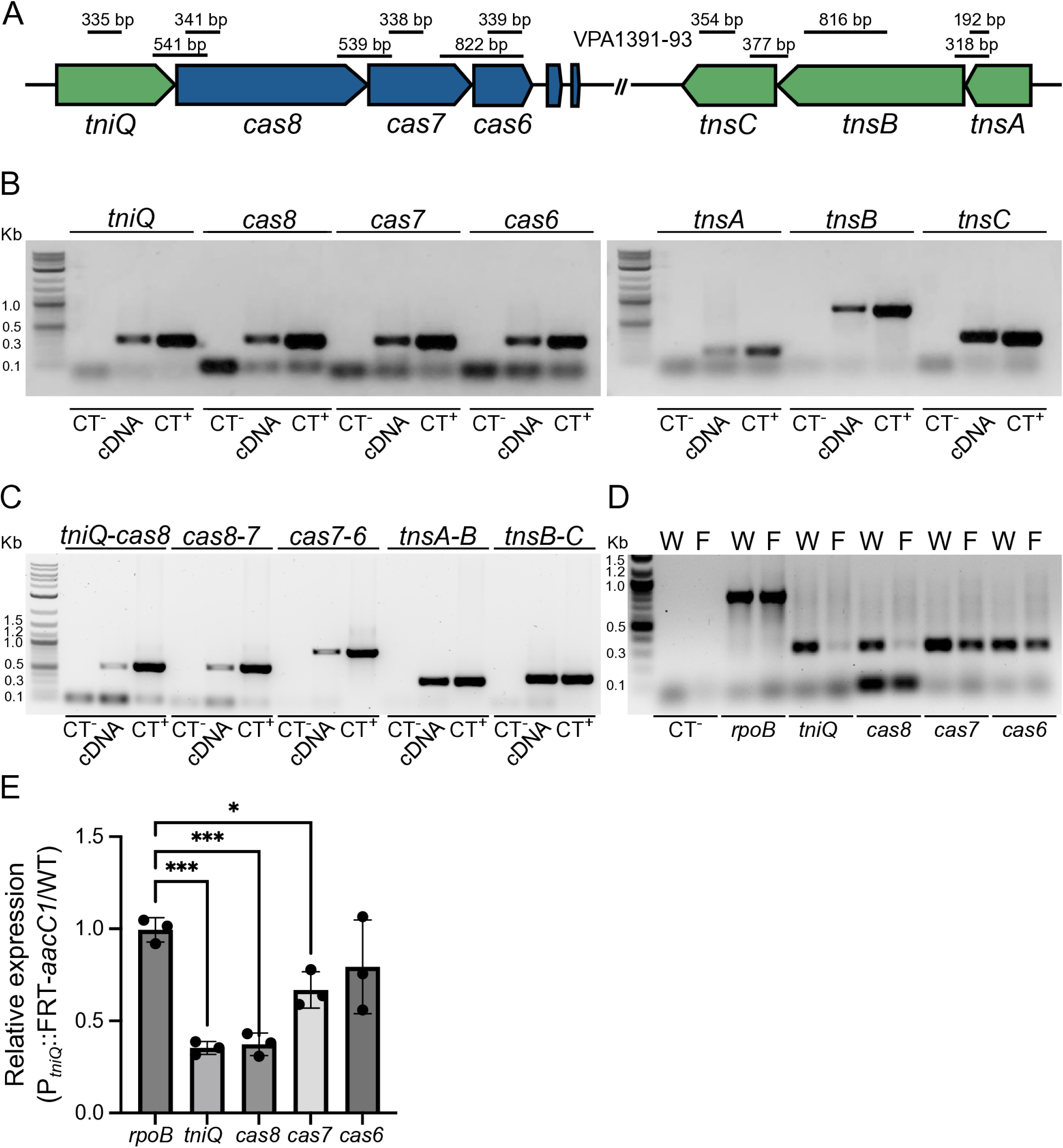
The genetic clusters *tniQ*-*cas876* and *tnsABC* are transcribed as polycistrons in *V. parahaemolyticus* RIMD2210633. A) Diagram illustrating the arrangement of the *tniQcas876* and *tnsABC* genetic clusters. The Tn7-like transposition associated genes are colored in green and the CRISPR/Cas associated genes are colored in blue. The lines shown above the genetic clusters indicate the location of the regions amplified by PCR, the size of the amplification product is also indicated. B, C) Image of the electrophoretic migration in an agarose gel of PCR products obtained with primer-pairs specific to B) monocistrons or C) polycistrons. D) Image of a representative RT-PCR analysis of the transcript abundance of *rpoB*, *tniQ*, *cas8*, *cas7* and *cas6* in the WT strain (W) or the P*_tniQ_*::FRT-*aacC1* mutant strain (F). Total RNA was used as template in negative controls (CT^-^), for panel D) a primer pair used to amplify a region of *rpoB* was used to assess contamination of total RNA with gDNA. Complementary DNA (cDNA) obtained by RT-PCR was used as template to detect the transcripts of interest. Genomic DNA from *V. parahaemolyticus* RIMD2210633 was used as template in positive controls (CT^+^). E) Bar graph representing the mean and standard deviation of the relative abundance (P*_tniQ_*::FRT*aacC1*/WT) of transcripts for *rpoB*, *tniQ*, *cas8*, *cas7* and *cas6* obtained through a semiquantitative RT-PCR analysis. Means were compared using an Ordinary one-way ANOVA test followed by a Dunnett’s test to compare each mean to the mean of the control *rpoB*. Mean differences with a P value ≤ 0.05 were interpreted as significant. * Adjusted P value ≤ 0.05, *** ≤ 0.001.

To further analyze the *tniQcas876* operon we compared the expression of these genes in a WT strain and a strain that has an insertion of an FRT-*aacC1* cassette immediately upstream of the first codon of *tniQ* (P*_tniQ_*::FRT-*aacC*1), most likely displacing its promoter. The *aacC1* gene, which encodes a Gentamicin 3-N-acetyltransferase, is positioned in a divergent orientation with respect to *tniQ*. We isolated total RNA from the WT strain and the P*_tniQ_*::FRT-*aacC*1 mutant strain in three independent experiments and did a semiquantitative RT-PCR analysis to evaluate the transcript abundance of *rpoB*, *tniQ, cas8, cas7* and *cas6*. RpoB is the β subunit of the RNA polymerase, and its expression should be the same in the WT strain and the P*_tniQ_*::FRT-*aacC*1 mutant strain. As expected, our results showed that the expression of *rpoB* was not altered when comparing the WT strain and the P*_tniQ_*::FRT-*aacC*1 mutant strain (Fig. 2DE). In contrast, the expression of *tniQ* and *cas8* decreased in the P*_tniQ_*::FRT-*aacC*1 mutant strain when compared to the expression in the WT strain (Fig. 2D). We observed a modest decrease in the abundance of *cas7* transcripts in the P*_tniQ_*::FRT-*aacC*1 mutant strain compared to the WT strain but could not detect significant changes in the expression of *cas6* with the resolution of our assay (Fig. 2DE). Regardless, our results suggest that the transcriptional activity of P*_tniQ_* mainly affects the expression of *tniQ* and *cas8* and that the *tniQcas876* genetic cluster could contain complex operons with internal promoters.

We next used the bacterial sigma70 promoter recognition program BPROM (28) to predict putative promoters upstream of *tniQ*, *cas8*, *cas7*, *cas6*, *tnsA*, *tnsB* and *tnsC*, the results are shown in Table 1. The BPROM tool predicted promoters upstream of all the genes analyzed, with different degrees of conservation with respect to the consensus of the −10, −10 extended and −35 boxes of sigma70 promoters from well characterized *E. coli* genes. The canonical A and T nucleotides located at positions −11 and −7 of the −10 box, respectively, are conserved in all predicted promoters. These nucleotides are important for the formation of the open complex and transcription initiation at the −10 box of sigma70 promoters (29).

**Table 1.**
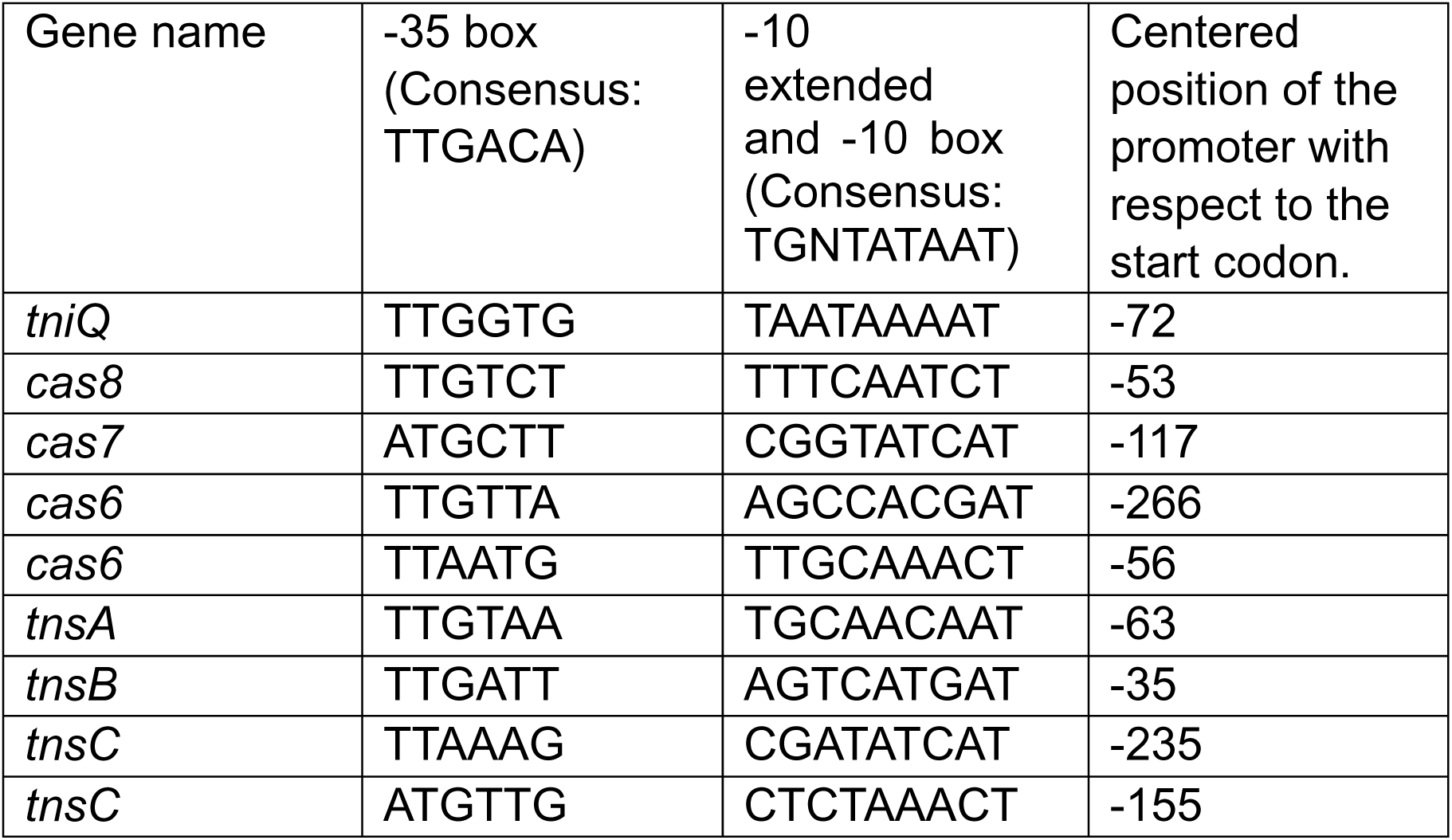
Predicted promoters within the putative *tniQcas876* and *tnsABC* operons.

To determine if the genes that conform the *V. parahaemolyticus* CRISPR/Cas-transposition system have independent promoters we generated transcriptional fusions to the reporter *luxCDABE* gene cluster using the sequence upstream of each structural gene as well as the CRISPR arrays (Fig. 3A). The *luxCDABE* cluster encodes a bacterial luciferase and its substrate, thus it can be used as a light-emitting genetic reporter. We used a strain that harbors the promoterless plasmid pBBRlux (P_empty_-*lux*) as negative control. We were able to identify promoters with activity significantly different from the negative control upstream of *tniQ*, *cas8*, *cas7*, *cas6*, the typical and atypical CRISPR arrays, *tnsA*, *tnsB* and *tnsC* (Fig. 3B). The transcriptional fusions with the upstream sequence from *cas7* and *cas8* showed low transcriptional activity with luminescence values close to those produced by the negative control, thus more work needs to be done to assess if these genes have independent promoters. The promoters upstream of *tniQ*, *tnsA* and *tnsB* showed similar levels of activity. The most active promoters were those of the two CRISPR arrays and *tnsC*.

**Figure 3.**
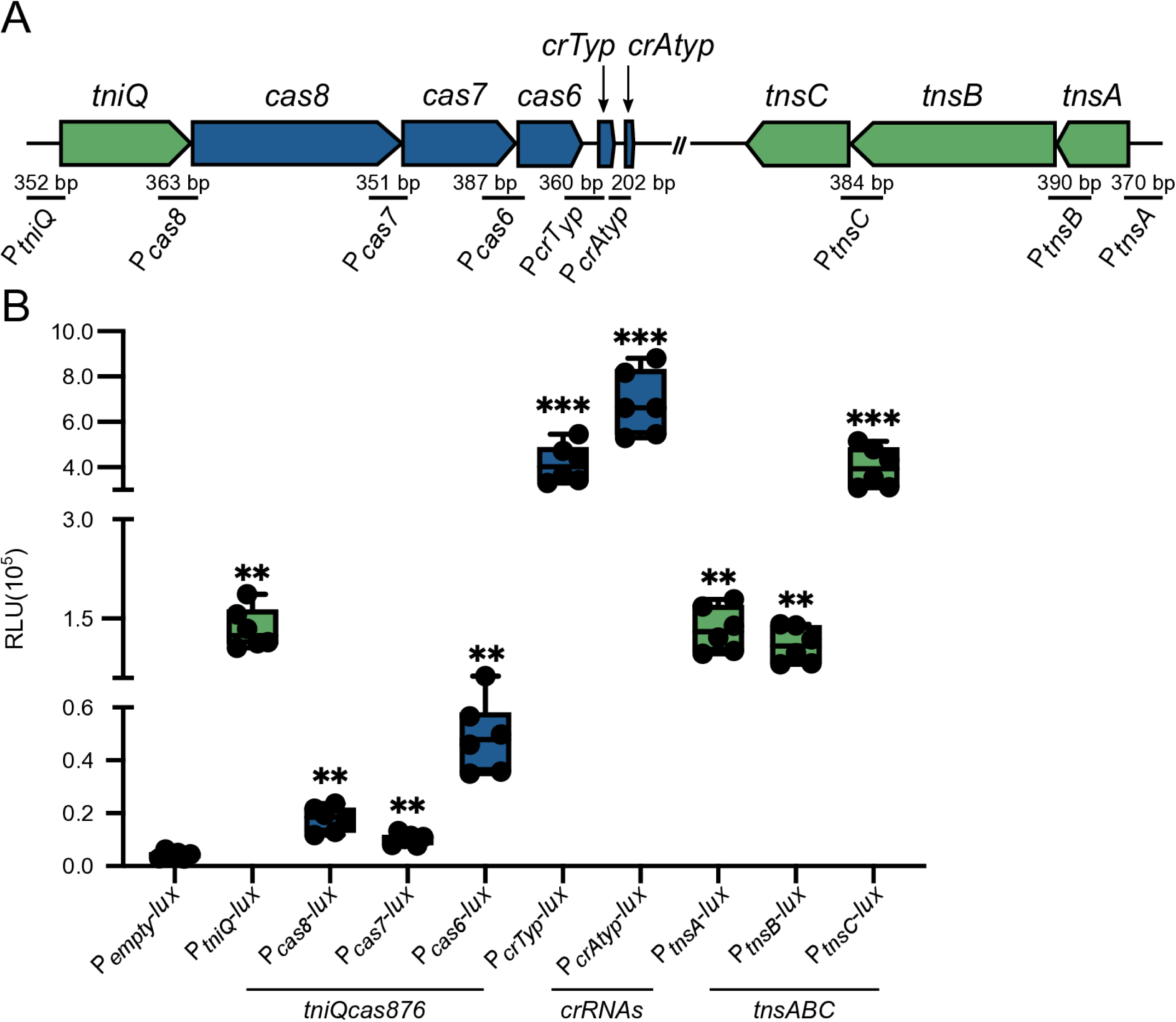
Multiple independent promoters drive transcription of genes associated with the *Vpa*CAST system. A) Diagram illustrating the genetic arrangement of the components of the *Vpa*CAST system, and the regions used to generate each transcriptional fusion with the *luxCDABE* gene cluster. B) Box plots represent the values of light production expressed as Relative Luminescence Units (RLU) obtained from 6 independent biological replicates of strains harboring the transcriptional fusions of interest. The label P_empty_ refers to the negative control of a strain harboring the promoterless plasmid pBBRlux. Means were compared using a Brown-Forsythe and Welch ANOVA test followed by a Dunnett’s T3 test to compare each mean to the mean of the negative control (P_empty_). Mean differences with a P value ≤ 0.05 were interpreted as significant. ** Adjusted P value ≤ 0.01, *** ≤ 0.001.

Our results suggest that the genes involved in CRISPR/Cas mediated transposition are transcriptionally active under standard laboratory growth conditions and that the *tniQcas876* and *tnsABC* genetic clusters are likely composed by complex operons. We found several independent promoters with variable levels of activity that could potentially provide tunable regulation and/or contribute to the overall accumulation levels of the components of the *Vpa*CAST system.

### The promoters of *tniQ*, the atypical CRISPR array, VPA1391 (*vpxR*) and VPA1392 are subject to transcriptional regulation in *V. parahaemolyticus*

Another fundamental question that we wanted to tackle in this work was whether the activity of the promoters that drive expression of the elements of the *Vpa*CAST system was subject to regulation. The product of VPA1391, which is encoded downstream of the atypical CRISPR array, is a putative transcriptional regulator of the Xre family which has previously been shown to positively regulate its own expression and negatively regulate the expression of the atypical CRISPR array from the *Vpa*CAST system. The evidence supporting these regulatory roles was obtained in the heterologous host *E. coli* and *in vitro* through biochemical experiments that demonstrated interaction of VPA1391 with a specific DNA sequence (23). We used a similar bioinformatic approach as previously reported to locate the VPA1391 binding site (VPA1391-box) upstream of VPA1391 and the Atypical CRISPR matrix (P*_crAtyp_*) (Fig 4A) (23). VPA1391 is encoded in the negative strand in a divergent orientation with respect to the gene VPA1392 which encodes a hypothetical protein (Fig. 4B). Since VPA1391 and VPA1392 share the same regulatory region, it is possible that both are regulated by the product of VPA1391 (Fig. 4B). The reported VPA1391-box overlaps with a putative −10 box element (TATAAT) from a putative promoter in both P*_crAtyp_* and P_VPA1392_ (Fig. 4B).

**Figure 4.**
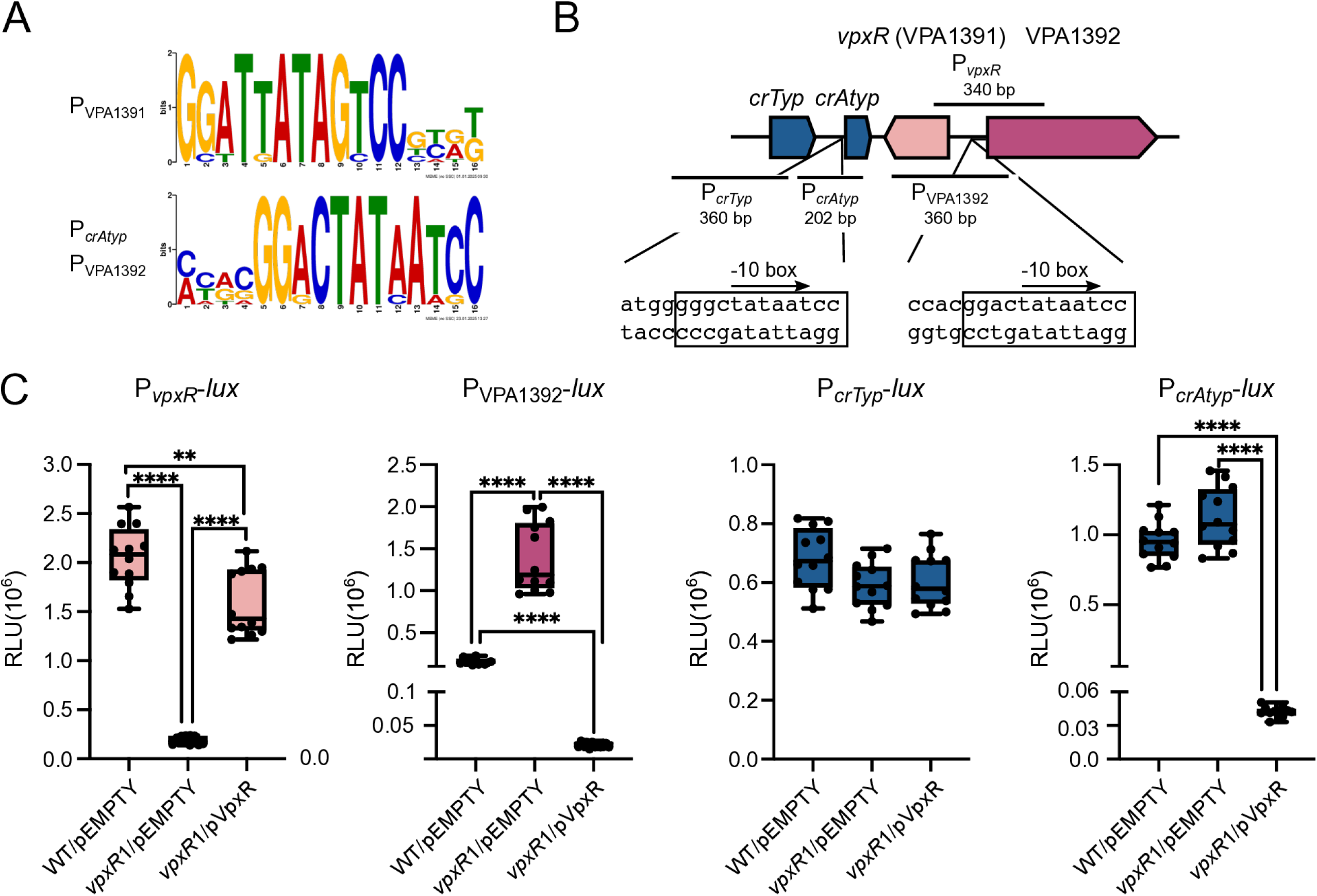
The VpxR (VPA1391) regulator positively regulates the P*vpxR* promoter and negatively regulates the P_VPA1392_ and P*_crAtyp_* promoters. A) Logo (Forward and reverse complement orientation) generated with frequency matrices of a conserved motif discovered by the Multiple Em for Motif Elicitation software. B) Diagram illustrating the arrangement of the CRISPR arrays and the *vpxR* and VPA1392 genes. The sequence of the VpxR-box and putative −10 boxes located upstream of the atypical CRISPR array and in the intergenic region between *vpxR* and VPA1392 is also shown. The putative VpxR binding-site is indicated with a rectangular box. C) Box plots represent the values of light production expressed as Relative Luminescence Units (RLU) obtained from 12 independent biological replicates from three independent experiments of strains harboring the transcriptional fusions of interest and either an empty overexpression plasmid (pEMPTY) or a plasmid that overexpresses *vpxR*. Means were compared using a Brown-Forsythe and Welch ANOVA test followed by a Dunnett’s T3 test for multiple comparisons. Mean differences with a P value ≤ 0.05 were interpreted as significant. **** P value ≤ 0.0001, ** P value ≤ 0.01.

To evaluate the relevance of the regulatory role of VPA1391, here named *Vpa*CAST Xre-like regulator (VpxR), in its native genetic background we first evaluated if it has an active promoter and if it is autoregulated. We generated a P*_vpxR_*-*lux* transcriptional fusion, an insertion mutation in *vpxR* using *Vch*INTEGRATE (*vpxR*1) and a plasmid that overexpresses *vpxR* (pVpxR) through the activity of an IPTG inducible promoter. Our results revealed that P*_vpxR_* is active in *V. parahaemolyticus* under standard laboratory growth conditions (Fig. 4C). The activity of P*_vpxR_* decreased significantly in a *vpxR*1 mutant strain compared to the WT strain, and the diminished activity of P*_vpxR_* in the *vpxR*1 mutant strain was complemented by expressing *vpxR in trans* (Fig. 4C). Together, these results strongly support that VpxR positively regulates its own transcription.

We next analyzed the effect of the absence or overexpression of *vpxR* on the activity of P_VPA1392_-*lux*, P*_crTyp_*-*lux* and P*_crAtyp_*-*lux*. The absence of *vpxR* resulted in approximately a 5-fold increase in the activity of P_VPA1392_ when compared to the activity in the WT strain (Fig. 4C). In contrast, the activity of P_VPA1392_ was approximately 24-fold higher in the WT strain compared to a strain that overproduces VpxR (Fig. 4C). These results strongly suggest that VpxR is a repressor of P_VPA1392_. Neither the absence nor the overexpression of *vpxR* affected the activity of P*_crTyp_* (Fig. 4C). This was somewhat expected since the regulatory region of P*_crTyp_* does not have a VpxR-box. Although the regulatory region of P*_crAtyp_* has a VpxR-box, we did not observe differences when comparing its activity in the WT strain and the *vpxR*1 mutant strain (Fig. 4C). However, the activity of P*_crAtyp_* in the WT strain was 17-fold higher than in a genetic background that overexpresses *vpxR* (Fig. 4C).

These results showed that VpxR is active in its native genetic background. The baseline production of VpxR under standard laboratory growth conditions is enough to activate P*_vpxR_* and repress P_VPA1392_. To observe the negative effect of VpxR on the activity of P*_crAtyp_* it was necessary to overexpress it from an IPTG inducible promoter.

While analyzing a bioinformatic prediction of potential regulatory targets of ToxR in *V. parahaemolyticus* we found that the regulatory region of *tniQ* had a consensus ToxR-box previously identified in ToxR-targets in *V. cholerae* (30) (Fig. 5A). This box is also present upstream of *leuO* on the reverse strand (Fig. 5A). LeuO, also named CalR in *V. parahaemolyticus*, is a transcriptional regulator of the LysR family whose transcription is positively regulated by ToxR in *V. cholerae* and in *V. parahaemolyticus* (31–34). Based on these observations we decided to evaluate if the activity of P*_tniQ_* was affected by the absence of ToxR. We used *Vch*INTEGRATE to generate an insertion mutation in *toxR* (*toxR*1). As a control we also analyzed the expression of a transcriptional fusion of P*_leuO_*-*luxCDABE*.

**Figure 5.**
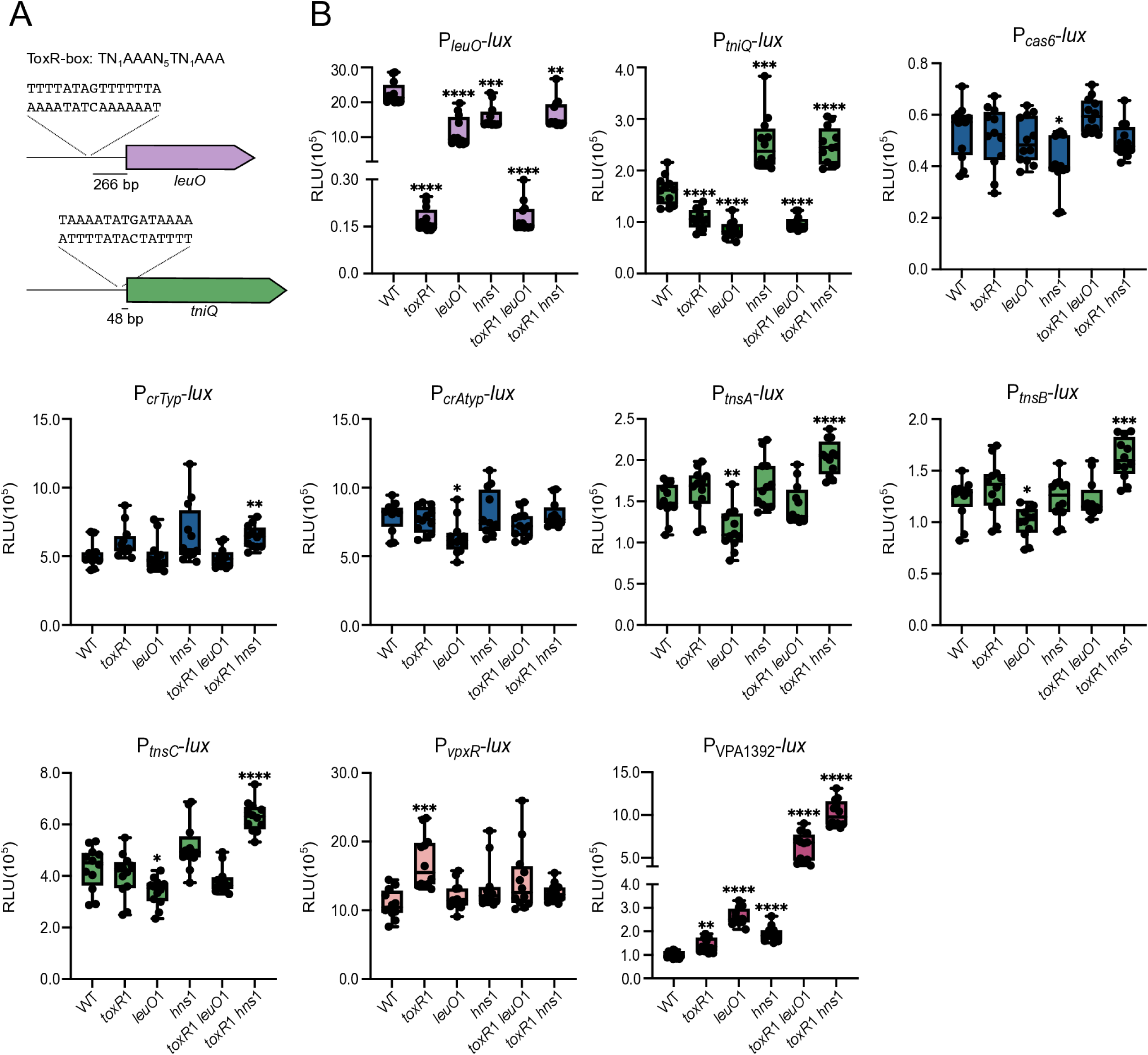
ToxR, LeuO and H-NS modulate the expression of the P*_tniQ_* and P_VPA1392_ promoters. A) Illustration depicting the sequence of the ToxR-box consensus and the putative ToxR-boxes located upstream of P*_leuO_* and P*_tniQ_* promoters. B) Box plots represent the values of light production expressed as Relative Luminescence Units (RLU) obtained from 12 independent biological replicates from three independent experiments of strains harboring the transcriptional fusions of interest. Means were compared using a BrownForsythe and Welch ANOVA test followed by a Dunnett’s T3 test for multiple comparisons. Mean differences with a P value ≤ 0.05 were interpreted as significant. **** P value ≤ 0.0001, *** P value ≤ 0.001, ** P value ≤ 0.01, * P value ≤ 0.05.

The activity of P*_leuO_* was 13-fold higher in the WT strain compared to that in the *toxR*1 mutant strain. The absence of ToxR had a modest negative effect on the activity of P*_tniQ_* showing a fold-change of 0.66 when compared to WT (Fig. 5B). ToxR and LeuO orthologues from multiple enterobacteria have been shown to antagonize the silencing effect of the histonelike nucleoid protein H-NS or coregulate H-NS targets (35–38). Based on this, we next evaluated if null mutations in *leuO* and *hns* affected the activity of P*_leuO_* and P*_tniQ_*. We used *Vch*INTEGRATE to generate single mutants in *leuO* (*leuO*1) and *hns* (*hns*1) and double mutants in *toxR* and *leuO* (*toxR*1 *leuO*1) and *toxR* and *hns* (*toxR*1 *hns*1). The activity of P*_leuO_* decreased 0.5-fold in the mutant strains *leuO*1, and *hns*1 compared to that in the WT strain (Fig. 5B). The activity of P*_leuO_* in the *toxR*1 *leuO*1 double mutant strain was the same as in the *toxR*1 single mutant strain, while its activity in the *toxR*1 *hns*1 double mutant strain was the same as in the *hns1* single mutant strain (Fig. 5B). These results support a role for ToxR as an antagonist of H-NS at the P*_leuO_* promoter. The activity of P*_tniQ_* showed a fold-change of 0.53 when comparing the *leuO1* mutant strain and the WT strain, and a 1.6 fold-change in activity when comparing the *hns1* mutant strain and the WT strain (Fig. 5B). On one hand, the activity of P*_tniQ_* in the *toxR1 leuO1* double mutant strain was comparable to the activity in the single *toxR*1 and *leuO*1 mutant strains (Fig. 5B). And on the other hand, the activity of P*_tniQ_* in the *toxR1 hns*1 double mutant strain was the same as in the *hns*1 single mutant strain. These results suggest that ToxR and/or LeuO could antagonize the modest silencing activity of H-NS at the P*_tniQ_* promoter.

We also evaluated the effect of the absence of ToxR, LeuO and H-NS on the activity of P*_cas6_* P*_crTyp_* P*_crAtyp_* P*_tnsA_* P*_tnsB_* P*_tnsC_* P_VPA1391_ and P_VPA1392_. The only promoter that showed fold-changes in activity above 1.5 when comparing the WT, single and double mutant strains was P_VPA1392_ (Fig. 5B). The activity of these promoter increased 2.6-fold in the *leuO* single mutant strain when compared to the WT strain. The combined absence of *leuO* and *toxR* or *toxR* and *hns* resulted in a 6-fold and 10-fold increase, respectively, in the activity of P_VPA1392_ when compared to the activity in the WT strain (Fig. 5B).

ToxR, LeuO and H-NS appear to modulate the expression of P*_tniQ_* but do not seem to be strong regulators of the *Vpa*CAST system under our experimental conditions. ToxR, LeuO and H-NS showed a significant silencing effect on the expression of VPA1392, however, it is not known if these open reading frame forms part of the *Vpa*CAST system.

### A synthetic construct expressing the *Vpa*CAST system promotes programmable insertion of a mini-CAST with different efficiency in *E. coli* compared to *V. parahaemolyticus*

Based on transposition assays carried in the heterologous host *E. coli*, it has been previously suggested that the elements involved in CRISPR/Cas mediated transposition from *V. parahaemolyticus* are functional (23, 24). Transposition mediated by the *Vpa*CAST system was found to be less efficient compared to transposition mediated by the *Vch*CAST system. To begin to investigate the potential roots of the differences in efficiency of these two CAST systems we compared the degree of conservation at the amino acid level between the proteins from their Cascade and Transposase complexes using EMBOSS

Needle (Table 2) (39). The TniQ, Cas8, Cas7 and Cas6 had % identities that go from 45% to 59%, while the TnsA, TnsB and TnsC proteins had % identities that go from 30% to 41%. We also analyzed the conservation of putative TnsB-boxes (16, 27) within the L-terminal and Rterminal sequences that delimit the *Vch*CAST and the *Vpa*CAST (Fig 1B). TnsB-boxes from the L-end were more conserved than the TnsB-boxes from the R-end (Fig 1B). The direct repeats of the typical CRISPR array from *Vch*CAST and the *Vpa*CAST systems were highly conserved, while the imperfect repeats from the atypical CRISPR arrays were not (Fig. 1C).

**Table 2.**
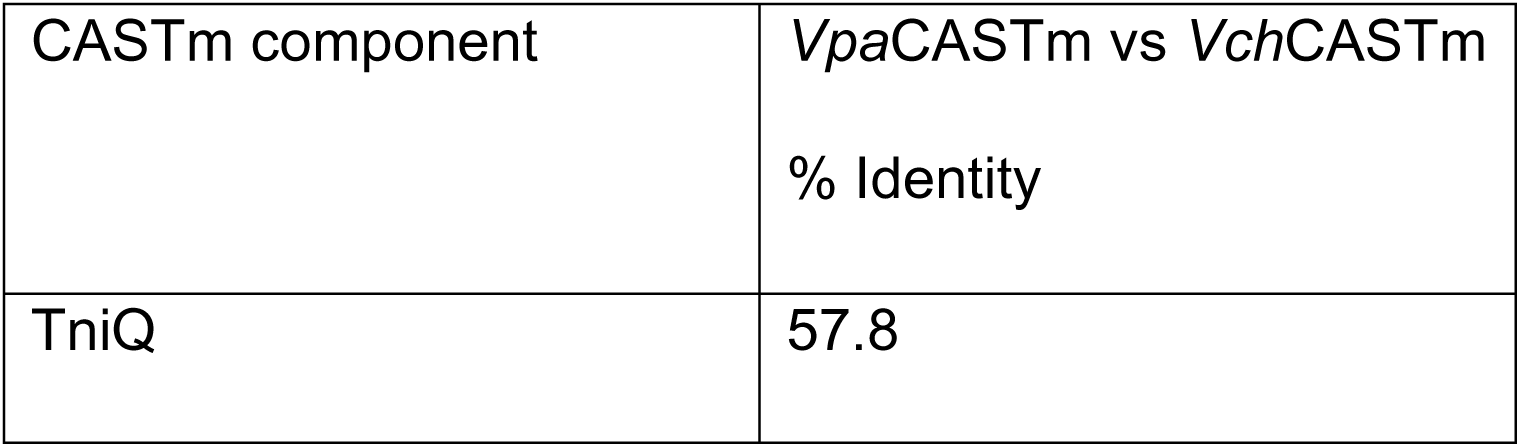
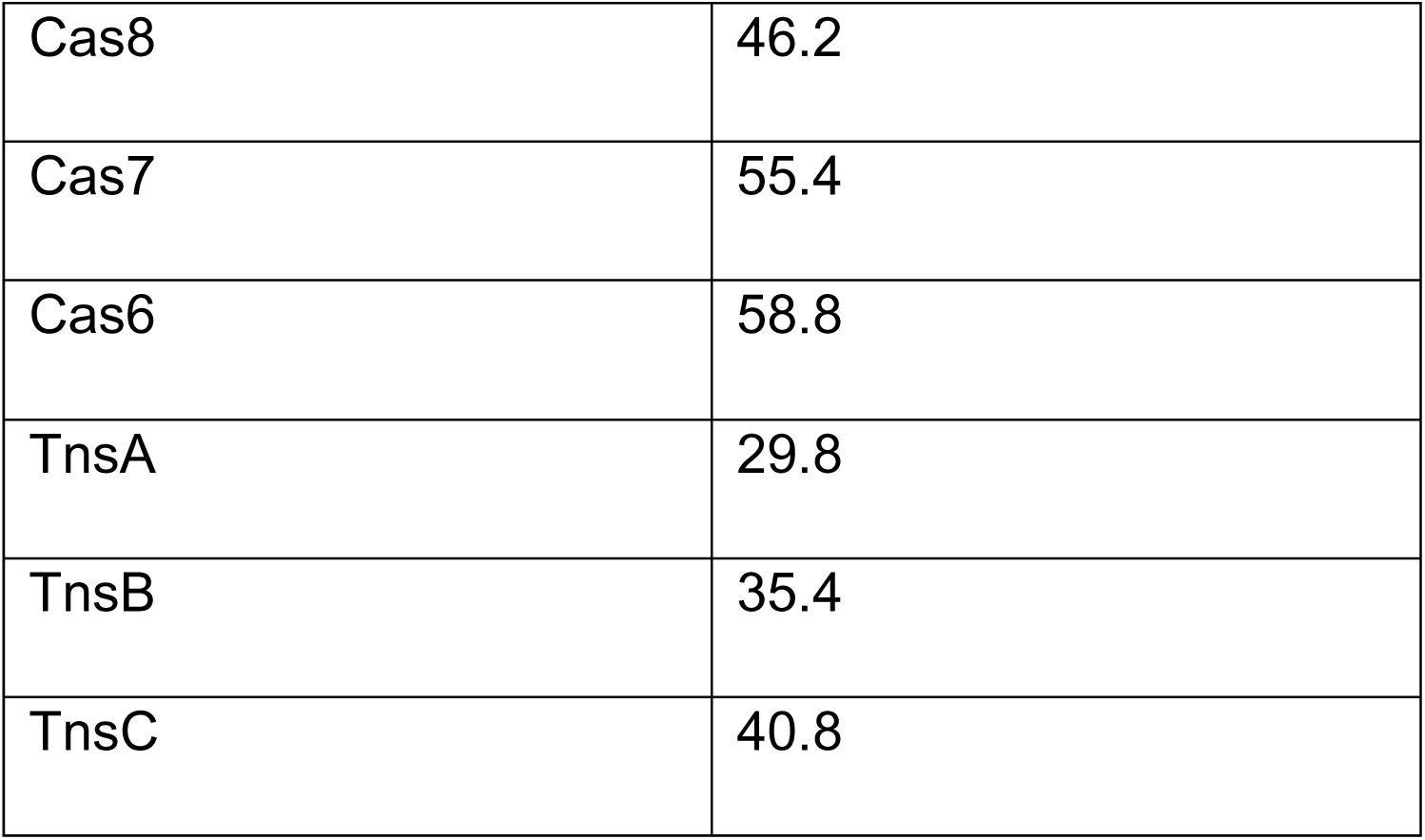
Pairwise comparison of amino acid identity between *Vpa*CASTm and *Vch*CASTm components.

It is not known if any of the above-mentioned differences are responsible for the variation in transposition efficiency mediated by these two CAST systems. We speculated that the *Vpa*CAST system may have adapted its function to the genetic context of *V*.

*parahaemolyticus,* and this had concomitantly caused it to lose activity in other hosts. To begin to tackle this hypothesis we analyzed *Vpa*CAST mediated transposition in *E. coli* and *V. parahaemolyticus*. To do this we followed a similar approach as previously reported for the generation of the gene editing tool INTEGRATE (27). This tool contains the elements required for CRISPR/Cas mediated transposition present in the transposon Tn6677 from *V. cholerae* HE-45 strain. These elements include the CRISPR/Cas-transposition machinery (CASTm) composed of TniQ, Cas8, Cas7, Cas6, TnsA, TnsB and TnsC, a CRISPR matrix composed of two direct perfect repeats and a programable spacer sequence, and a mini CRISPR associated transposon (mini-CAST) flanked by right and left-terminal sequences containing a non-symmetric arrangement of recognition sites for TnsB (Fig. 1B). All these elements are assembled on a pBBR1MCS-2 plasmid backbone that is named pSL1142 and we refer to here as *Vch*INTEGRATE. To generate a *V. parahaemolyticus* based INTEGRATE plasmid (*Vpa*INTEGRATE), we replaced all the components corresponding to the CASTm, the CRISPR matrix and mini-CAST in pSL1142 for sequences corresponding to the counterpart elements located in VpaI-7 in chromosome 2 of *V. parahaemolyticus* RIMD2210633 (Fig. 6A).

**Figure 6.**
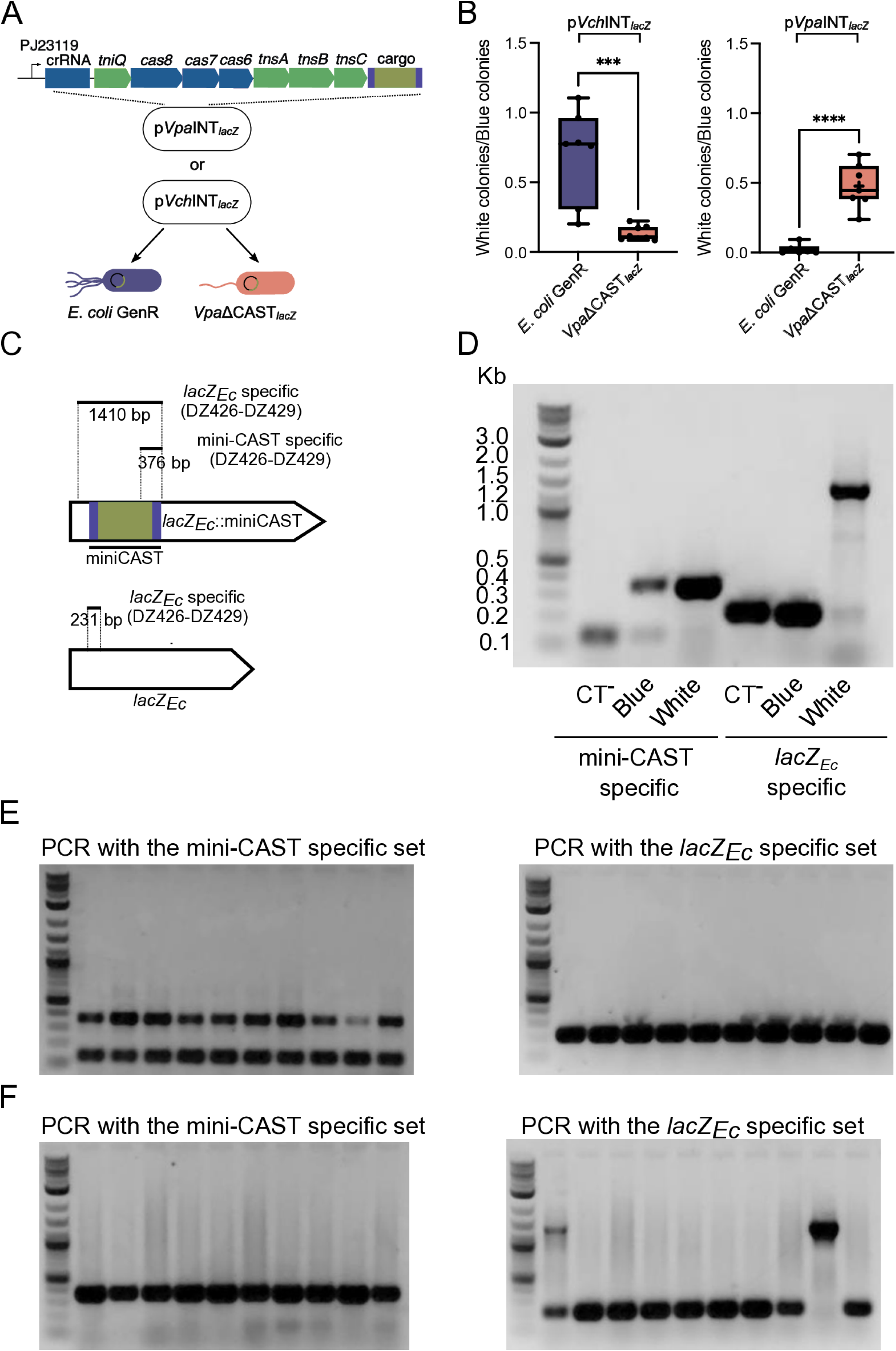
The CRISPR/Cas-transposition system from *V. parahaemolyticus* promotes transposition in *E. coli* and *V. parahaemolyticus*. A) Diagram illustrating the screening assay used to analyze transposition events in the *lacZ_Ec_* allele from *E. coli* GenR and *Vpa*ΔCAST*_lacZ_* strains harboring either the p*Vch*INT*_lacZ_* or p*Vpa*INT*_lac_*_Z_ plasmid. B) Box plots of the proportion of white colonies over blue colonies from *E. coli* GenR and *Vpa*ΔCAST*_lacZ_* strains harboring either the p*Vch*INT*_lacZ_* or p*Vpa*INT*_lac_*_Z_ plasmid grown in selection plates with X-gal and IPTG. Experiments were done at least six times. The differences between means were analyzed with an Unpaired t-test. Mean differences with a P value ≤ 0.05 were interpreted as significant. **** indicate a P value ≤ 0.0001. ** indicate a P value ≤ 0.01. C) Diagram of the regions amplified by PCR to identify transposition events and intact *lacZ_Ec_* copies. The solid lines represent the amplified regions and the region corresponding to the mini-CAST. D) Image of the electrophoretic migration in an agarose gel of PCR products stained with ethidium bromide (EtBr) and obtained using as template the lysate of a blue colony from a *V. parahaemolyticus* strain harboring p*Vpa*INT (CT^-^), or blue and white colonies from a *V. parahaemolyticus* strain harboring p*Vpa*INT*_lacZ_*. E, F) Image of the electrophoretic migration in an agarose gel of PCR products stained with EtBr and obtained using as template the lysate of 10 randomly selected colonies of E) *E. coli* or F) *V. parahaemolyticus* strains harboring p*Vpa*INT*_lacZ_*. All gels were loaded in the first lane with the Quick-Load Purple 1 kb Plus DNA Ladder from New England Biolabs. The relevant size range of the ladder is indicated on the left gel from panel B.

It has been documented that a CAST insertion event generates immunity towards new CAST transposition events (16, 27). Since the *V. parahaemolyticus* RIMD2210633 genome has all the elements of a CAST, we removed these elements to generate a genetic background that lacks the CASTm, CRISPR arrays and the L and R-terminal sequences that are recognized by TnsB. We will refer to this genetic background as ΔCAST from here on. To facilitate the detection of transposition events through genetic screenings we generated a reporter strain with a ΔCAST, *lacZ_Ec_* genotype (*Vpa*ΔCAST*_lacZ_*). In contrast to the WT strain and the ΔCAST mutant strain, *Vpa*ΔCAST*_lacZ_* produces a functional LacZ enzyme (from *E. coli*) that is capable of degrading X-gal and thus allows to screen for colonies with βGalactosidase activity based on blue versus white colony-coloration (Fig. 6A). We reprogrammed the CRISPR matrix from the *Vpa*INTEGRATE plasmid (p*Vpa*INT) with a 32 base pair sequence that encodes a crRNA that is complementary to *lacZ_Ec_* (p*Vpa*INT*_lacZ_*) (Fig. 6A). As a positive control we also reprogrammed the *Vch*INTEGRATE plasmid (p*Vch*INT) with the same *lacZ_Ec_*-targeting crRNA to generate p*Vch*INT*_lacZ_*. As negative controls we mobilized through conjugation the non-targeted plasmid p*Vch*INT and p*Vpa*INT to a derivative of *E. coli* BL21 DE3 that is gentamicin resistant (*E. coli* GenR) or to *Vpa*ΔCAST*_lacZ_*. All the transconjugant colonies showed a blue coloration when grown in the presence of X-gal (Data not shown). We next evaluated if p*Vch*INT*_lacZ_* and p*Vpa*INT*_lacZ_* were able to promote transposition of the mini-CAST inside *lacZ_Ec_* in *E. coli* GenR and in

*Vpa*ΔCAST*_lacZ_*. The proportion of white vs blue colonies in transconjugants of *E. coli* GenR and *Vpa*ΔCAST*_lacZ_* with p*Vch*INT*_lacZ_* was in average 0.7 and 0.13 respectively (Fig. 6B). This would suggest that under the conditions tested the *Vch*INTEGRATE system was more efficient in promoting transposition in *E. coli* GenR than in *Vpa*ΔCAST*_lacZ_*. In contrast, the proportion of white vs blue colonies in transconjugants of *E. coli* GenR and *Vpa*ΔCAST*_lacZ_* with p*Vpa*INT*_lacZ_* was in average 0.02 and 0.5 respectively (Fig. 6B). This would suggest that the *Vpa*INTEGRATE system is more efficient in promoting transposition in *Vpa*ΔCAST*_lacZ_* than in *E. coli* GenR. These results support previous observations that showed that the *Vch*CAST system was more efficient than the *Vpa*CAST system in promoting transposition in *E. coli*. Furthermore, our data suggest that host factors might affect the transposition efficiency mediated by CAST systems but more work needs to be done to fully demonstrate that this is the case.

To further evaluate transposition events, we used an end point PCR-based approach that required two primer pairs. The first pair (DZ432-DZ429) here referred to as the mini-CAST specific set, only allows detection of transposition events (Fig. 6C). The second pair (DZ426-DZ429) here referred to as the *lacZ_Ec_* specific set, amplifies either a short sequence of 231 bp corresponding to an intact *lacZ_Ec_* allele or a 1410 bp sequence corresponding to an insertion event of the mini-CAST in *lacZ_Ec_*, (Fig. 6C).

We first used these primer sets in *Vpa*ΔCAST*_lacZ_* strains that received either p*Vpa*INT (negative control) or p*Vpa*INT*_lacZ_* and that were grown in screening plates containing X-gal and IPTG. When we used the lysate of a blue colony from the negative control strain as template for amplification, as expected, the mini-CAST specific primer set did not amplify the 376 bp amplicon corresponding to transposition events. We did observe a smaller unspecific band with a size between 100 and 200 bp (Fig. 6D). When we used a lysate of a blue colony of strain *Vpa*ΔCAST*_lacZ_*/p*Vpa*INT*_lacZ_* as template for amplification with the miniCAST specific primer set we obtained both the specific 376 bp amplicon corresponding to transposition events and the unspecific amplicon observed in the negative control (Fig. 6D). This would suggest that even in blue colonies from strains harboring p*Vpa*INT*_lacZ_* there might be cells with transposition events. When we used the lysate of a white colony of strain *Vpa*ΔCAST*_lacZ_*/p*Vpa*INT*_lacZ_* as template for PCR with the mini-CAST specific primer set we only obtained the 376 bp amplicon that is specific for transposition events in *lacZ_Ec_* (Fig. 6D). When we used the *lacZ_Ec_* specific primer set we were able to identify intact copies of *lacZ_Ec_* in the blue colonies of the negative control and the *Vpa*ΔCAST*_lacZ_*/p*Vpa*INT*_lacZ_* strain, and interrupted copies of *lacZ_Ec_* only in the white colony from the *Vpa*ΔCAST*_lacZ_*/p*Vpa*INT*_lacZ_* strain (Fig. 6D). We also observed a faint band that could correspond to intact copies of *lacZ_Ec_* in the white colony, suggesting that it may still have cells without a transposition event (Fig. 6D). Since the blue coloration of a colony suggests a majority of *lacZ*^+^ cells and the white coloration a majority of *lacZ*^-^ cells, our results suggest that the mini-CASTspecific primer set can detect rare transposition events within a colony, while the *lacZ_Ec_* specific primer set can only detect transposition events when they occur at a high proportion within a colony.

We next used the two primer sets to analyze transposition events in 10 randomly selected colonies from *E. coli* GenR/p*Vpa*INT*_lacZ_* and *Vpa*ΔCAST*_lacZ_*/p*Vpa*INT*_lacZ_* strains grown in the absence of X-gal. The 10 colonies analyzed from strain *E. coli* GenR/p*Vpa*INT*_lacZ_* had transposition events that were revealed when we used the mini-CAST specific primer set but not with the *lacZ_Ec_* specific primer set (Fig. 2E). The 10 colonies analyzed from strain *Vpa*ΔCAST*_lacZ_*/p*Vpa*INT*_lacZ_* had transposition events that were revealed when we used the mini-CAST specific primer set, and two of these colonies had transposition events that were also revealed when we used the *lacZ_Ec_* specific primer set. Thus, the PCR results shown in Fig. 2E and F support our observation that transposition efficiency promoted by the *Vpa*INTEGRATE system is likely higher in the *V. parahaemolyticus* background compared to the *E. coli* background.

### The promoters that control the expression of the *Vpa*CAST system are more active in *V. parahaemolyticus* than in *E. coli*

The p*Vpa*INT plasmid used in the experiments from Fig. 6 contains several of the internal promoters that we found to be transcriptionally active. We speculated that a possible explanation for the differences in efficiency of the *Vpa*INTEGRATE plasmid in promoting transposition of a mini-CAST in an *E. coli* vs *V. parahaemolyticus* genetic background could be that these internal promoters are less active in *E. coli* than in *V*. *parahaemolyticus*. We compared the transcriptional activity in *V. parahaemolyticus* and *E. coli* of the active promoters identified in Fig. 3B, and the synthetic promoter P_J23119_ used to drive expression of the INTEGRATE tool. All the promoters from the genes of the *Vpa*CAST system were less active in *E. coli* than in *V. parahaemolyticus* (Fig. 7). In contrast, P_J23119_ showed a 1.6-foldincrease in activity in *E. coli* compared to *V. parahaemolyticus* (Fig. 7). The P*_cas6_* promoter showed almost no activity in *E. coli*. The rest of the promoters showed between 2- and 4fold increased activity in *V. parahaemolyticus* compared to *E. coli*, except for the promoter P*_tnsB_* which showed approximately 19-fold higher expression in *V. parahaemolyticus* compared to *E. coli* (Fig. 7).

**Figure 7.**
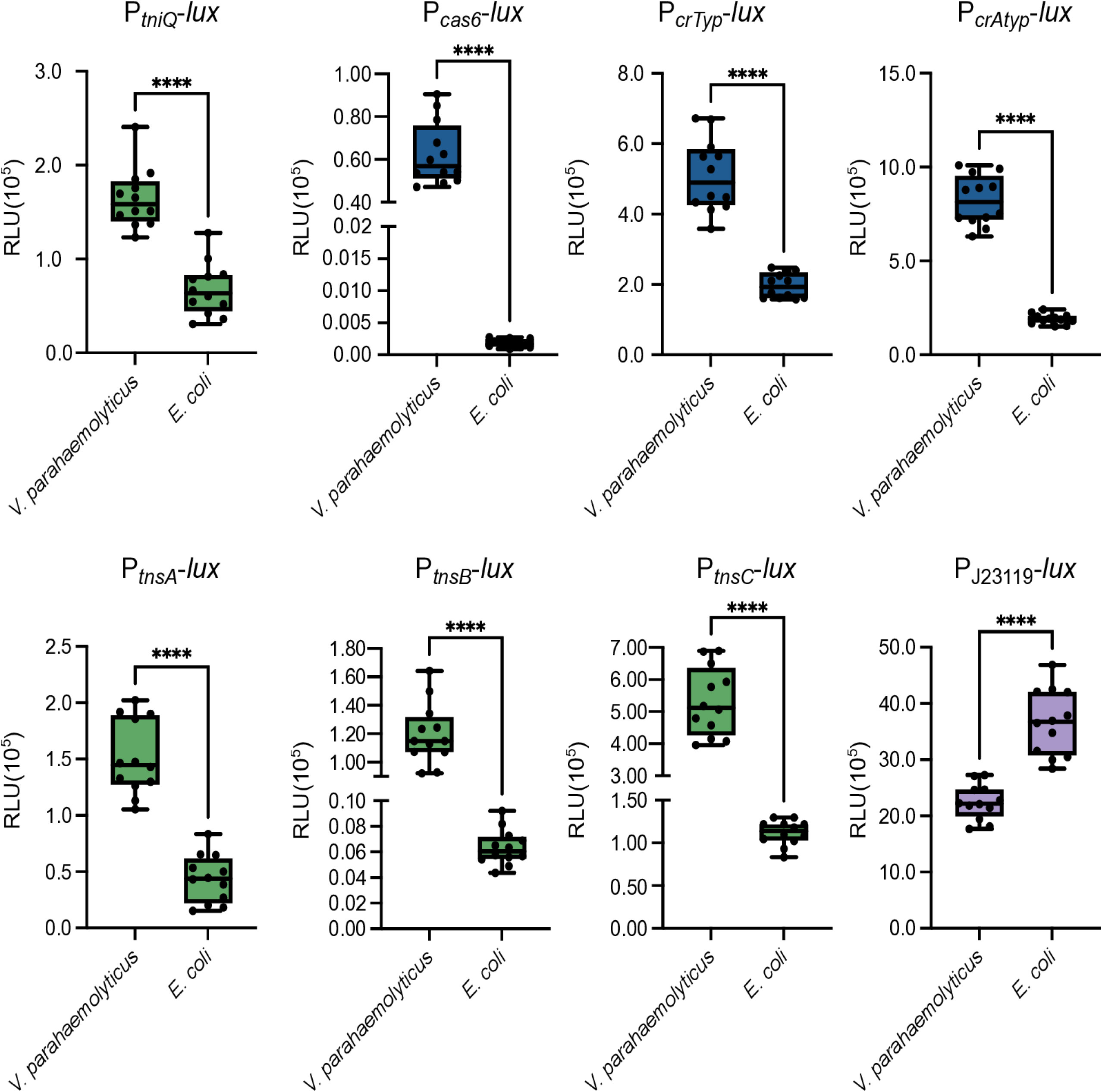
The promoters that drive transcription of genes associated with the *Vpa*CAST system are more active in *V. parahaemolyticus* than in *E. coli*. Box plots represent the values of light production expressed as Relative Luminescence Units (RLU) obtained from 12 independent biological replicates of *V. parahaemolyticus* and *E. coli* strains harboring the transcriptional fusions of interest. The data was obtained from three independent experiments. Pairwise comparisons of means were done using an Unpaired t-test. Mean differences with a P value ≤ 0.05 were interpreted as significant.. **** P value ≤ 0.0001.

With these observations alone we cannot conclusively affirm that the differences in transposition efficiency mediated by the *Vpa*CAST system in *V. parahaemolyticus* vs *E. coli* are due to changes in expression levels of its components, but we consider this should be a factor to be consider in future studies.

## Discussion

The recently discovered CAST systems are an example of how the coevolution of mobile genetic elements (MGEs) and bacterial defense systems can result in the emergence of biological innovations. Although our understanding of the transposition capacity, structure and regulation of different CASTs has progressively improved, we still have limited insight on how these MGEs operate in their native hosts (15, 24, 27, 40–45). Evaluating the behavior of CASTs in their native context could improve our understanding of their function; and learning how they are regulated and their impact on cell physiology could unveil biological principles of the mechanisms that govern the coevolution of these MGEs and their hosts. Here we report the first genetic characterization of the CAST I-F3a system from *V. parahaemolyticus* RIMD2210633, which is part of the VpaI-7 pathogenicity island and is possibly responsible for disseminating virulence factors (T3SS2 and TDH) between *Vibrio* strains in the environment. A key piece of information that could suggest that the *Vpa*CAST system is relevant for the physiology of *V. parahaemolyticus* RIMD2210633 is that of the transcriptional activity of its components. An *in silico* analysis of the nucleotidic and amino acid sequence of the *Vpa*CAST components suggests that they are somewhat conserved when compared to the components of the *Vch*CAST system. No clear indications of frame shifts or other deleterious mutations within the structural genes associated with the *Vpa*CAST system were found (24). However, mutations or sequence determinants that could affect their transcription are harder to identify by visual inspection. Here we showed that the *tniQcas876* and *tnsABC* genes are transcribed under standard laboratory growth conditions, they form polycistronic transcripts in *V. parahaemolyticus*, and have several internal promoters. The biological significance of these internal promoters needs to be further analyzed within their genetic context to determine how meaningful they are to the expression of the *Vpa*CAST system.

We found that the regulator VpxR (VPA1391) is expressed and controls the activity of the P*_vpxR_* and P*_vpa1392_* promoters in *V. parahaemolyticus*. Under the conditions tested, the overexpression but not the baseline production of VpxR repressed the transcriptional activity of the P*_crAtyp_* promoter. Finally, we showed that the regulators ToxR, LeuO, and H-NS modulate the expression of *tniQ*. We did not observe a clear effect of these regulators on the transcription of other genes from the VPA1387-VPA1396 region except for VPA1392. We cannot rule out that analyzing these regulatory regions outside of their native genetic context could result in topological changes that mask the potential effects of VpxR, ToxR, LeuO and H-NS on their transcription. More work needs to be done to determine the physiological relevance of the regulation of these genes by the above-mentioned regulators and perhaps additional transcriptional regulators.

The implication of having an actively transcribed *Vpa*CAST system are still unclear. We still do not know if the transcription levels of the *Vpa*CAST system observed in this study are sufficient to promote transposition, and we are currently working in tackling this question. It would seem wasteful to have unregulated production of an MGE that can only be mobilized to specific targets that might not be always available inside the cell. However, a constitutive expression of the components of the *Vpa*CAST system is perhaps necessary for an efficient transposition towards plasmids acquired through conjugation or natural competency. The typical CRISPR array located in the VpaI-7 island has two spacers, one of which targets plasmids such as pL300 that can be hosted by *V. alginolyticus* strains (25, 26). It will be of great importance to determine if the constitutive expression of the typical crRNA from the *Vpa*CAST system is sufficient to promote transposition into plasmids of ecological relevance. The regulation of the *Vpa*CAST system at the transcriptional level and beyond, needs to be further studied, as well as the relevance of its baseline level of expression for yet unidentified physiological roles.

Previous studies performed in the heterologous host *E. coli* showed that the Xre regulator encoded in VpaI-7 here referred to as VpxR acts as a repressor of the activity of the P*_crAtyp_* promoter (23), in addition to autoregulating its own expression through a mechanism similar to that of the repressor protein CI from phage lambda (46–48). The strong repression exerted by VpxR against P*_crAtyp_* led the authors to propose that CAST mobility is negatively regulated at the transcriptional level when the CAST has been established inside a host (23). However, our findings suggest that the atypical CRISPR array and *vpxR* are expressed under standard laboratory growth conditions, and that the expression of the former is not affected by the absence of the latter. The repressive nature of VpxR over the activity of the P*_crAty_* promoter was only observed when the regulator was artificially overexpressed. We did observe the autoregulation of *vpxR* and found that it negatively regulates the divergent gene VPA1392 which encodes a hypothetical protein. The VpxR-box found upstream of P*_crAty_* and P_VPA1392_ has some differences that could possibly affect the strength of the interaction of VpxR with each of these two regulatory regions. Our group is currently focused on the characterization of these regulatory regions and on analyzing if the role of VpxR goes beyond the control of the expression of the atypical CRISPR array from VpaI-7.

Another key observation that casts some doubts on the functionality of the *Vpa*CAST system is its reduced ability to promote transposition of synthetic mini-CASTs in *E. coli* when compared to the ability of the *Vch*CAST system (24). This differences in functionality could be due to several factors. An interesting observation is that of the apparent separate evolutionary trajectories of the *tniQcas876* and *tnsABC* operons even among close related bacteria (13, 24). Apparently, the components of the *tniQcas876* operon are subject to evolutionary forces that result in higher conservation at the DNA and protein level, while the components of the *tnsABC* system appear to be more divergent. To our knowledge there has not been a clear association with divergency and changes in functionality other than target categorization to avoid cross-talk among related transposition systems, but they could potentially explain why some CAST systems are more efficient than others in promoting transposition (23, 24). The divergence in TnsA, TnsB and TnsC is accompanied also by divergency in the TnsB-boxes that are recognized by the transposase complex (24, 45). While the three TnsB-boxes present in the L-terminal sequence of the *Vpa*CAST closely resemble the consensus CA[A/C]CCATA[A/T] [A/G]NTGATA[T/A][T/C][T/G] of TnsB-boxes present in *Vch*CAST systems, the TnsB-boxes from the R-terminal sequence significantly diverged (24, 45). It is not exactly known how these deviations from the consensus in the TnsB-boxes affect the affinity of TnsB for the R-terminal sequence. As mentioned above, it has been suggested that these sequence changes were selected to avoid cross-talk between CAST systems, it would be interesting to evaluate in the future if loss of efficiency can be compensated by the emergence of novel *in trans* or *cis* regulatory mechanisms. In this study we provided evidence that suggests that the transposition efficiency of the *Vpa*CAST system is higher in its native host, *V. parahaemolyticus*, than in *E. coli*. The ramifications of this observation need to be studied further to better understand the conditions and factors that can lead to differences in transposition efficiency in different genetic backgrounds. We observed that the internal promoters within the *tniQcas876* and *tnsABC* operons were more active in *V. parahaemolyticus* than in *E. coli*, so perhaps they accumulate to higher concentrations in the former. The threshold of *Vpa*CASTm accumulation needed to achieve maximal transposition efficiency is unknow and will be important to define in the future. Beyond the transcriptional-regulation explanation, it is tempting to speculate that ‘host factors’ could intervene to facilitate the transposition process in genetic backgrounds where this MGEs have evolved. Such host factors have been reported and include ACP and L29 that favor Tn7-transposition in *E. coli* (49), IHF and ClpX, enhance transposition efficiency of the *Vch*CAST system (45, 50), and the S15 ribosomal subunit that does so for the CRISPR associated transposition system of *Scytonema hofmannii* (*Sho*CAST system) (51, 52). The identification of potential ‘*Vibrio* integration factors’ would be of great relevance to understand the reach and limitations of CASTs in shaping the evolution of *Vibrio* species. Furthermore, the identification of such factors could also help to broaden the host range of *Vch*CAST and *Vpa*CAST-based engineering tools.

CRISPR associated transposition appears to have played an important role in the evolution of a variety of bacterial species, the fact that the components of the CASTm encoded in several genomes retain functionality opens the question of what their active contribution to horizontal gene transfer in the environment is. Here we reported our initial findings on the characterization of a CAST system that could have played an important role in the emergence of pandemic attributes in *V. parahaemolyticus*. Based on the potential that these MGEs have to propagate virulence factors, we consider of great relevance to continue the study of the factors that limit or favor their capacity for horizontal gene transfer. A better understanding of the functionality of the CAST systems in their native host could inform us how they might have shaped or could actively be shaping the evolution of human pathogens such as *V. parahaemolyticus*.

## Material and methods

### Strains, plasmids, and growth conditions

The strains and plasmids used in this study are described in Table 3. All strains were grown in Lysogeny Broth (LB) (1% tryptone, 0.5% yeast extract, and 1% NaCl) or on LB agar (LB with 1.5% bacteriological agar). Strains derived from *V. parahaemolyticus* RIMD2210633 were grown at 30°C and *E. coli* strains were grown at 37°C. Strains and plasmids were selected by adding antibiotics to the growth media at the following concentrations: streptomycin (Str) at 200 µg/mL, kanamycin (Kan) at 30 µg/mL, gentamicin (Gen) at 15 µg/mL, and chloramphenicol (Chl) at 20 µg/mL for *E. coli* and 5 µg/mL for *V. parahaemolyticus*.

**Table 3.**
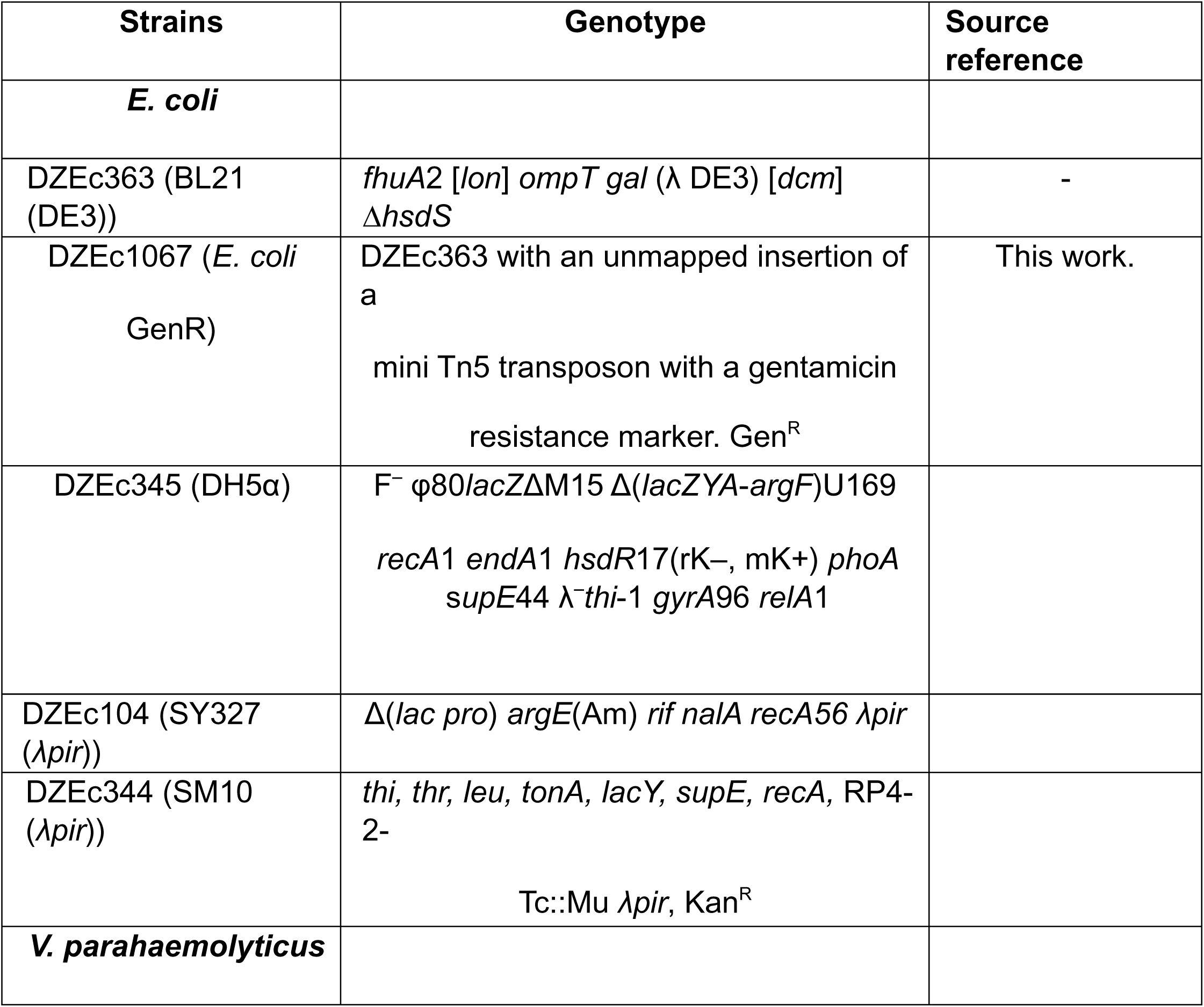
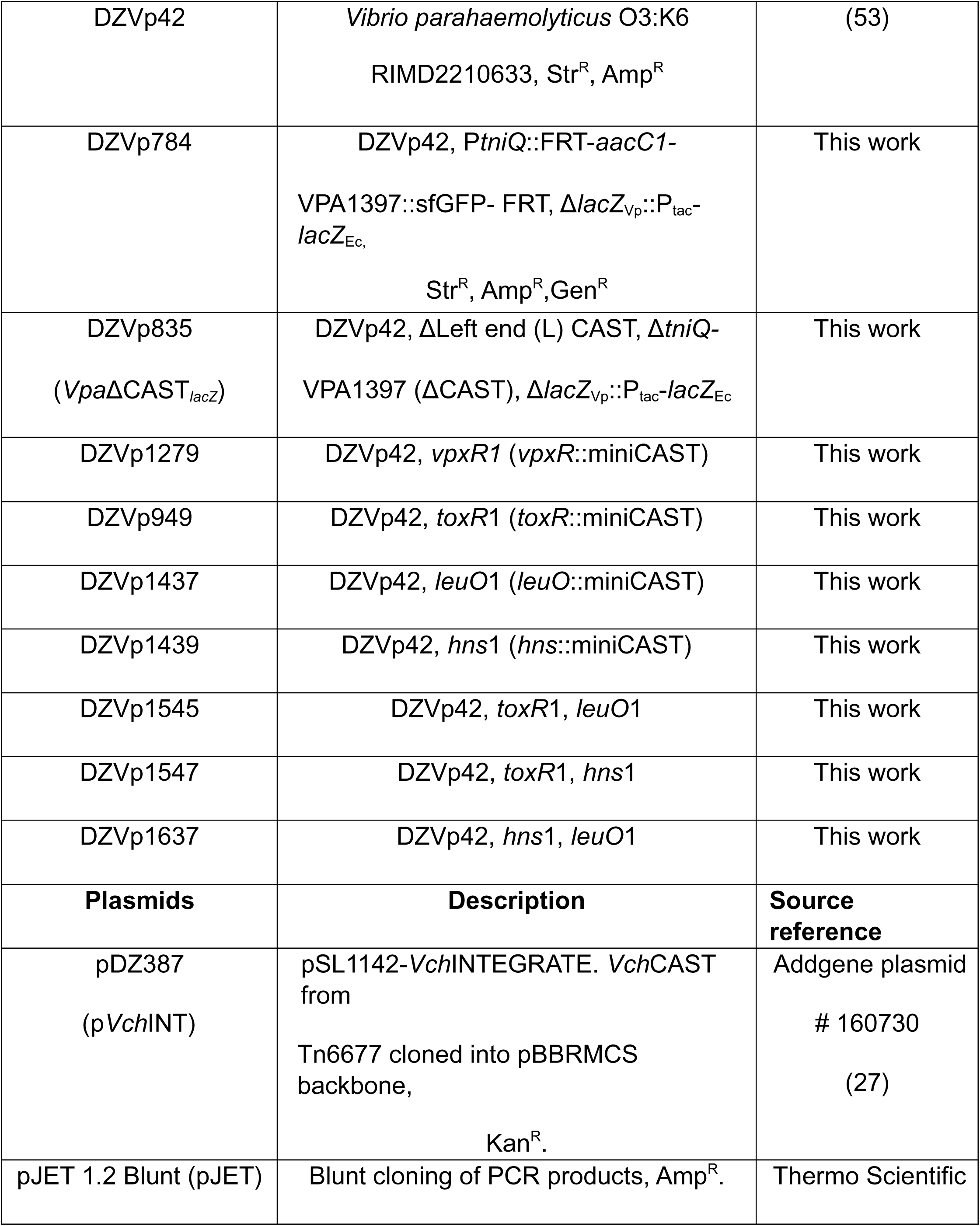
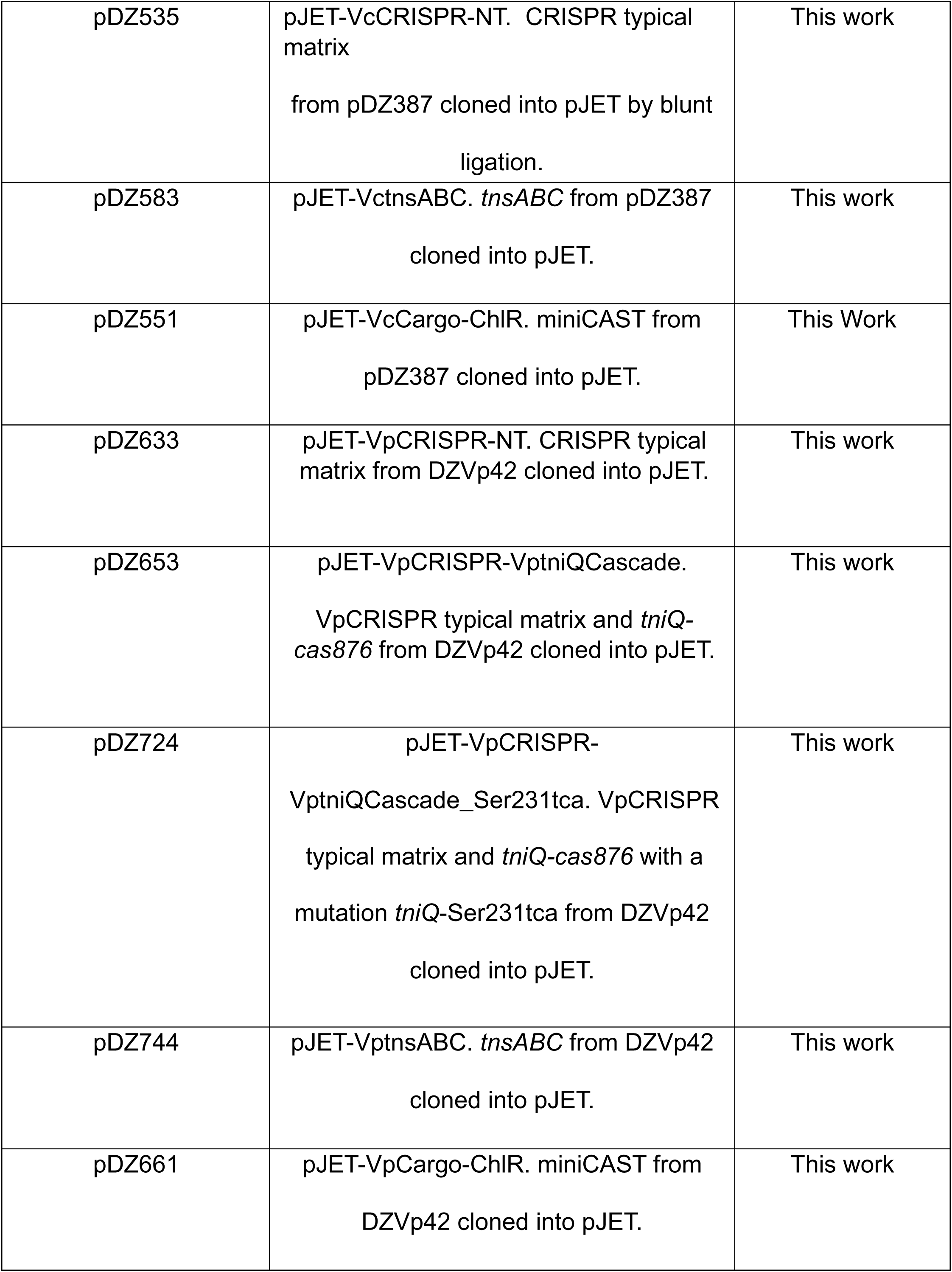
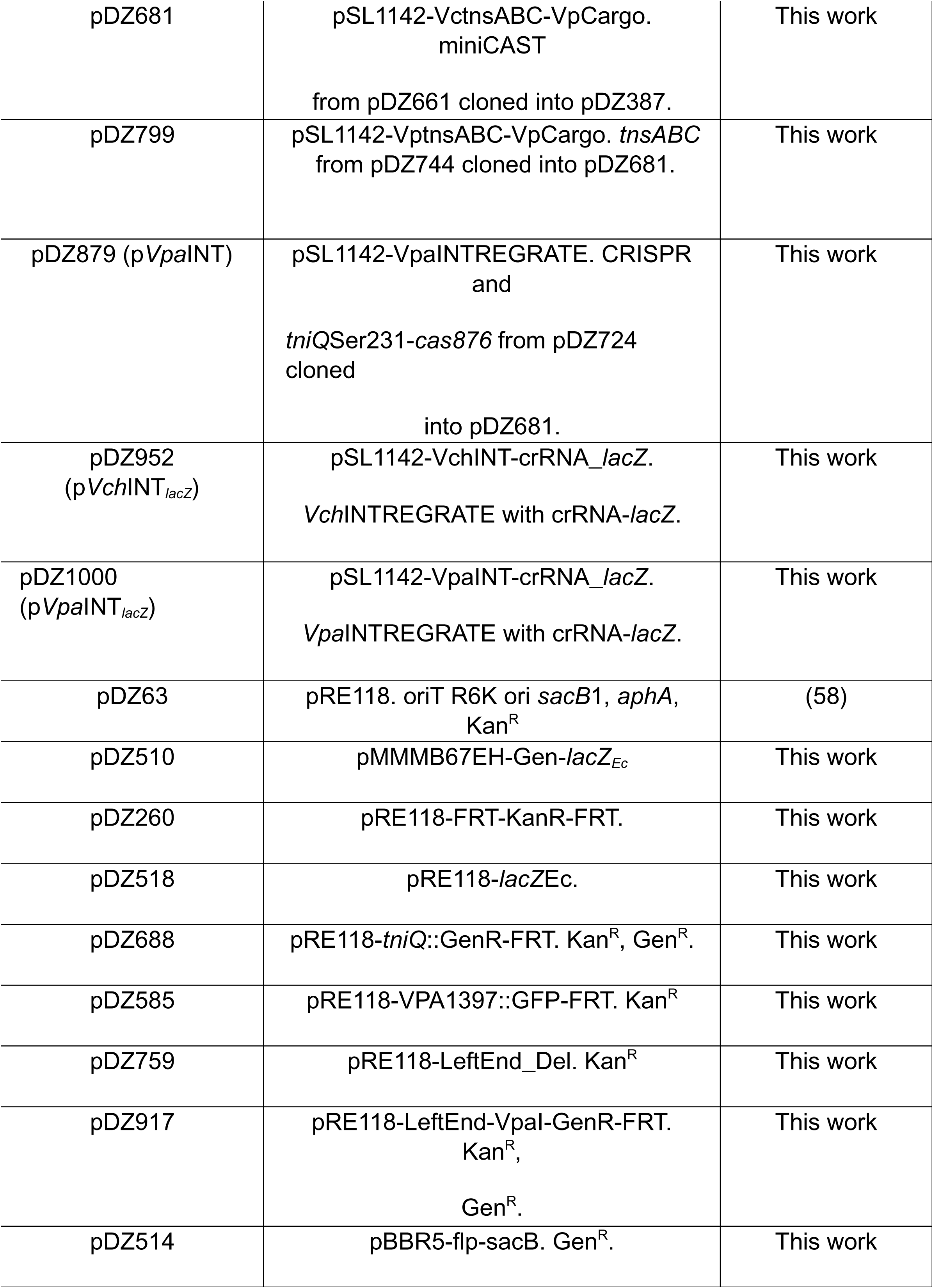
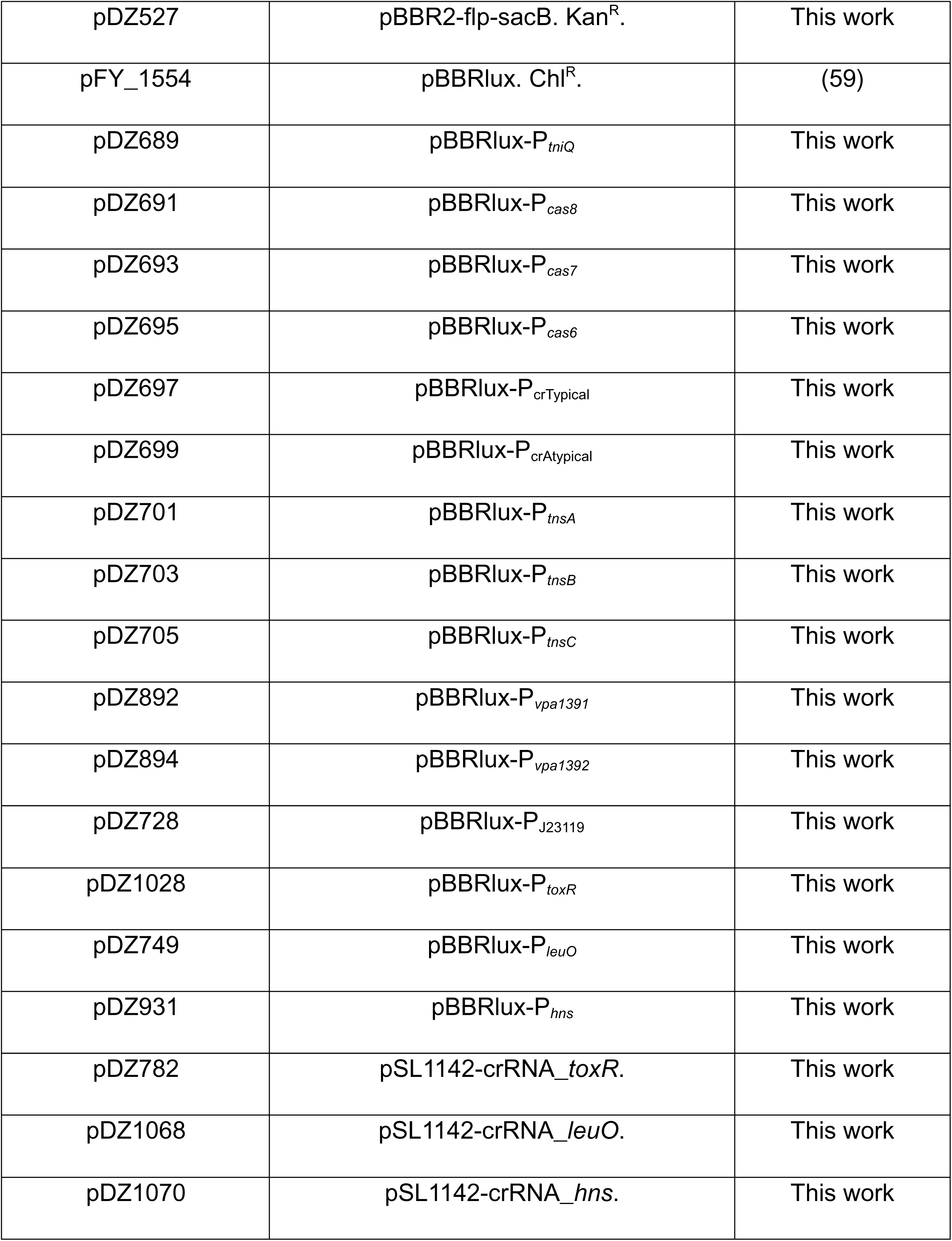
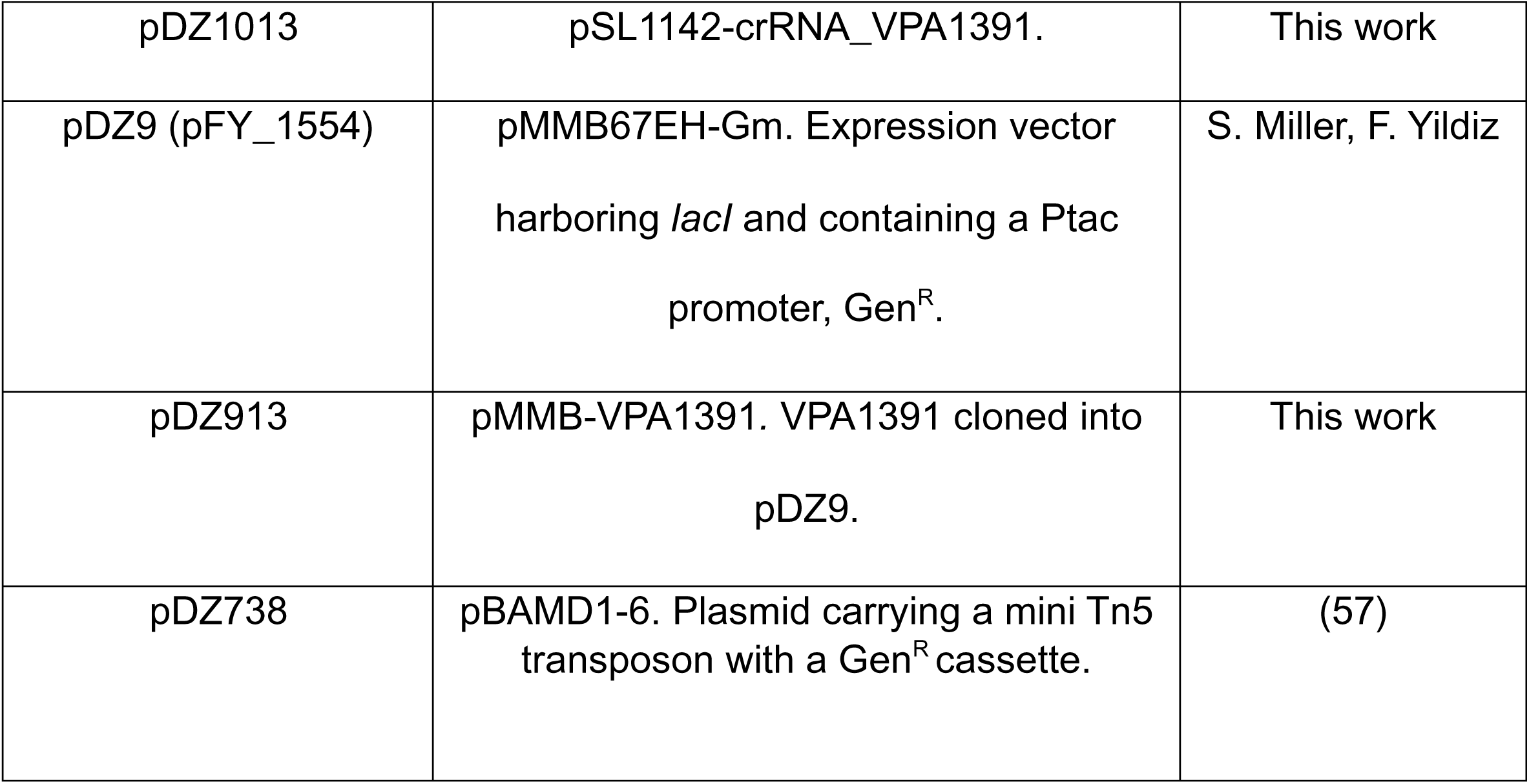
Strains and plasmids used in this study.

### RNA purification and cDNA synthesis

RNA was extracted from cells from strains DZVp42 (WT) and DZVp784 (P*_tniQ_*::FRT-*aacC1*) grown to exponential phase in LB broth at 30°C with agitation, using the commercial kit Direct-zol RNA MiniPrep from Zymo Research following the instructions of the manufacturer. The total RNA was concentrated and treated with DNase I using the commercial kit RNA Clean and Concentrator-5 from Zymo Research. The complete removal of DNA from the extractions was verified through PCR.

The DNAase I-treated RNA was used as template to synthesize cDNA with the commercial reagent LunaScript® RT SuperMix following the manufacturer instructions.

The RT-PCR semiquantitative analysis was done from three independent experiments. The densitometric analysis of DNA bands stained with Ethidium Bromide was done using the Volume tools of the analysis software Image Lab from BioRad.

### Generation of genetic constructs

The primers used for the generation of genetic constructs are described in Table 4. Amplicons used for assembling genetic constructs were produced by Polymerase chain reaction (PCR) with the high-fidelity DNA polymerase Q5 from New England Biolabs. PCR purification and plasmid isolation was done using the DNA Clean & Concentrator-5 and Zyppy Plasmid Miniprep Kits from Zymo Research, respectively. Sanger-sequencing was used to validate the fidelity of the genetic constructs generated for this study.

**Table 4.**
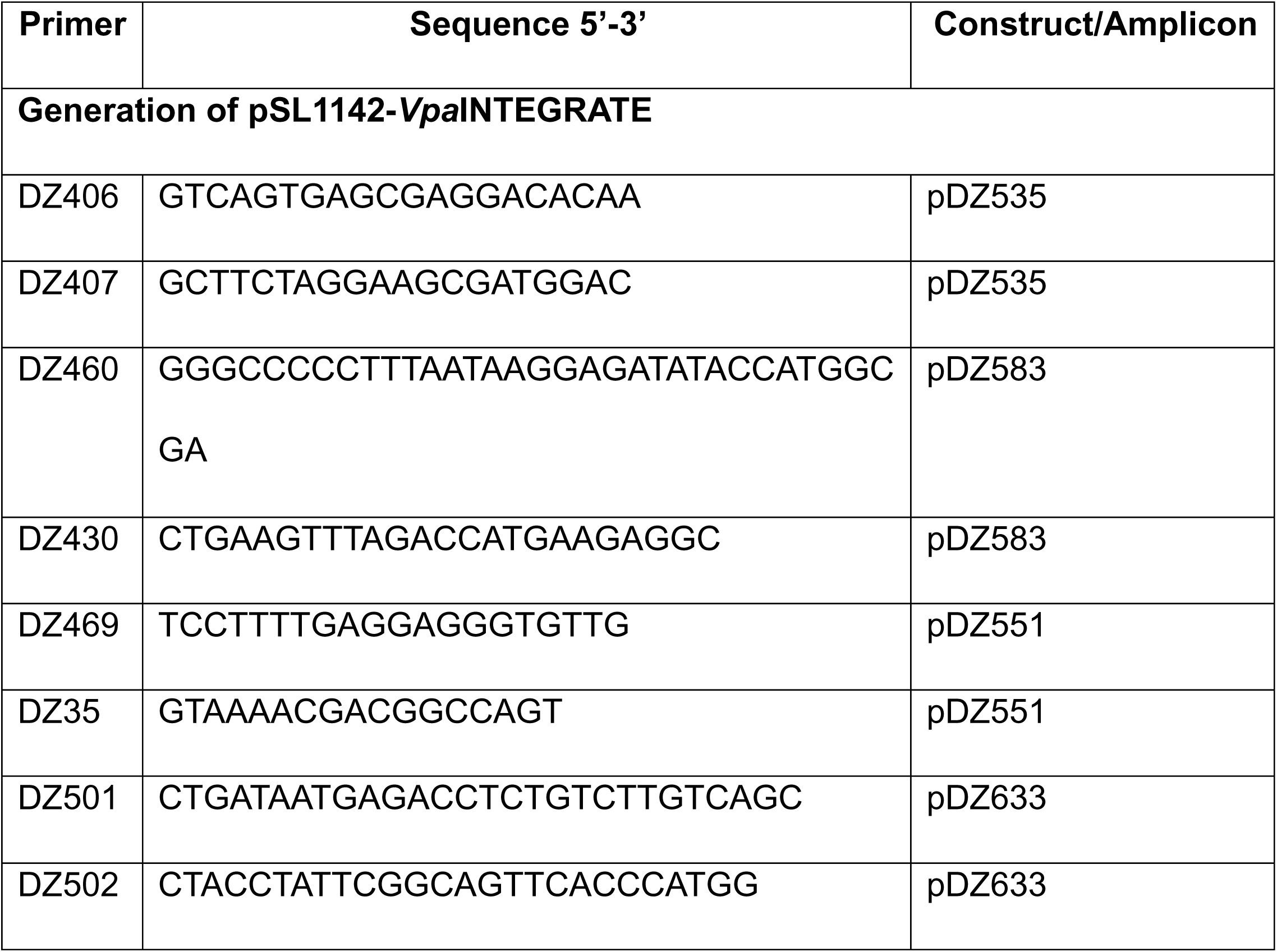
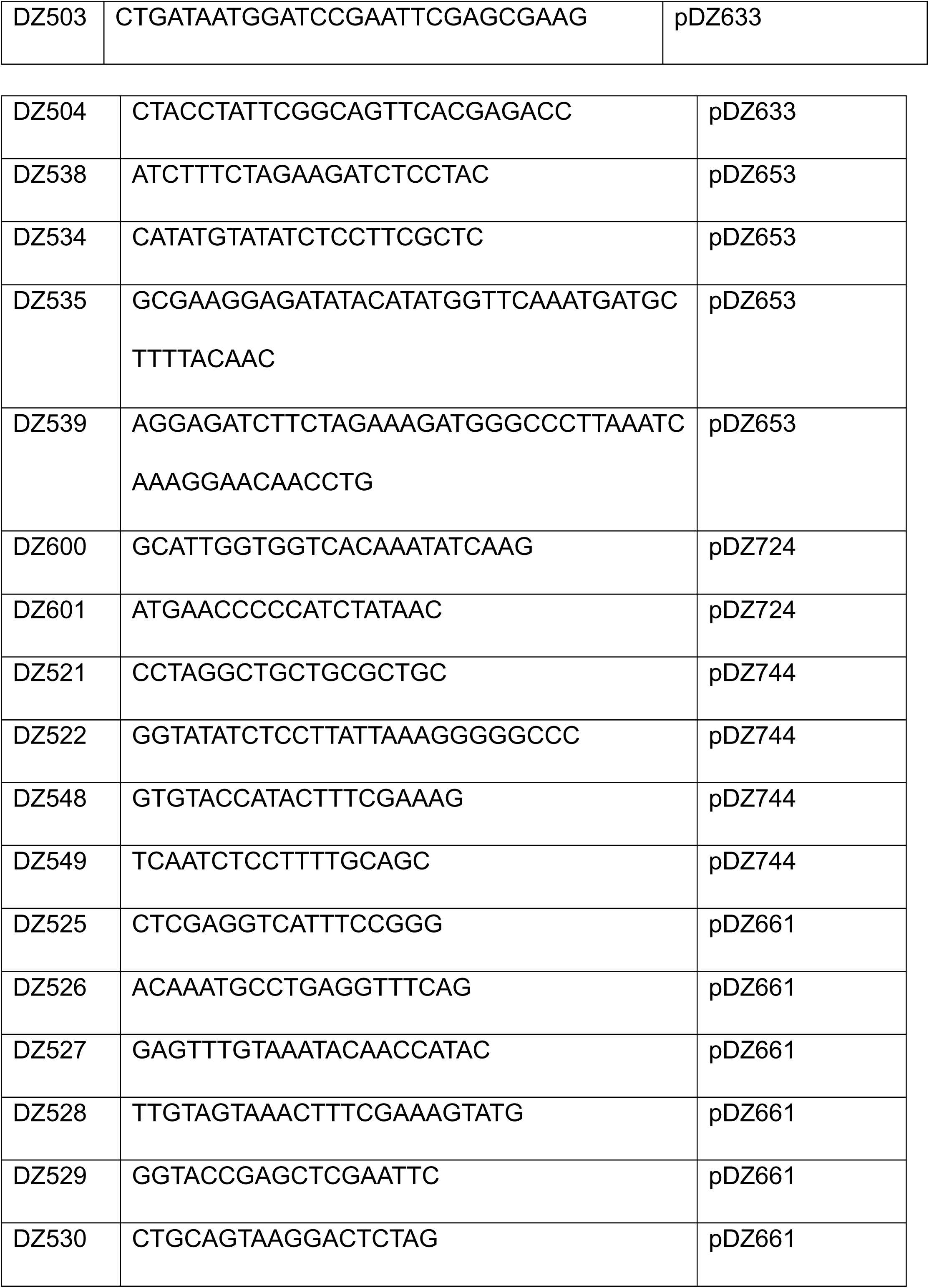
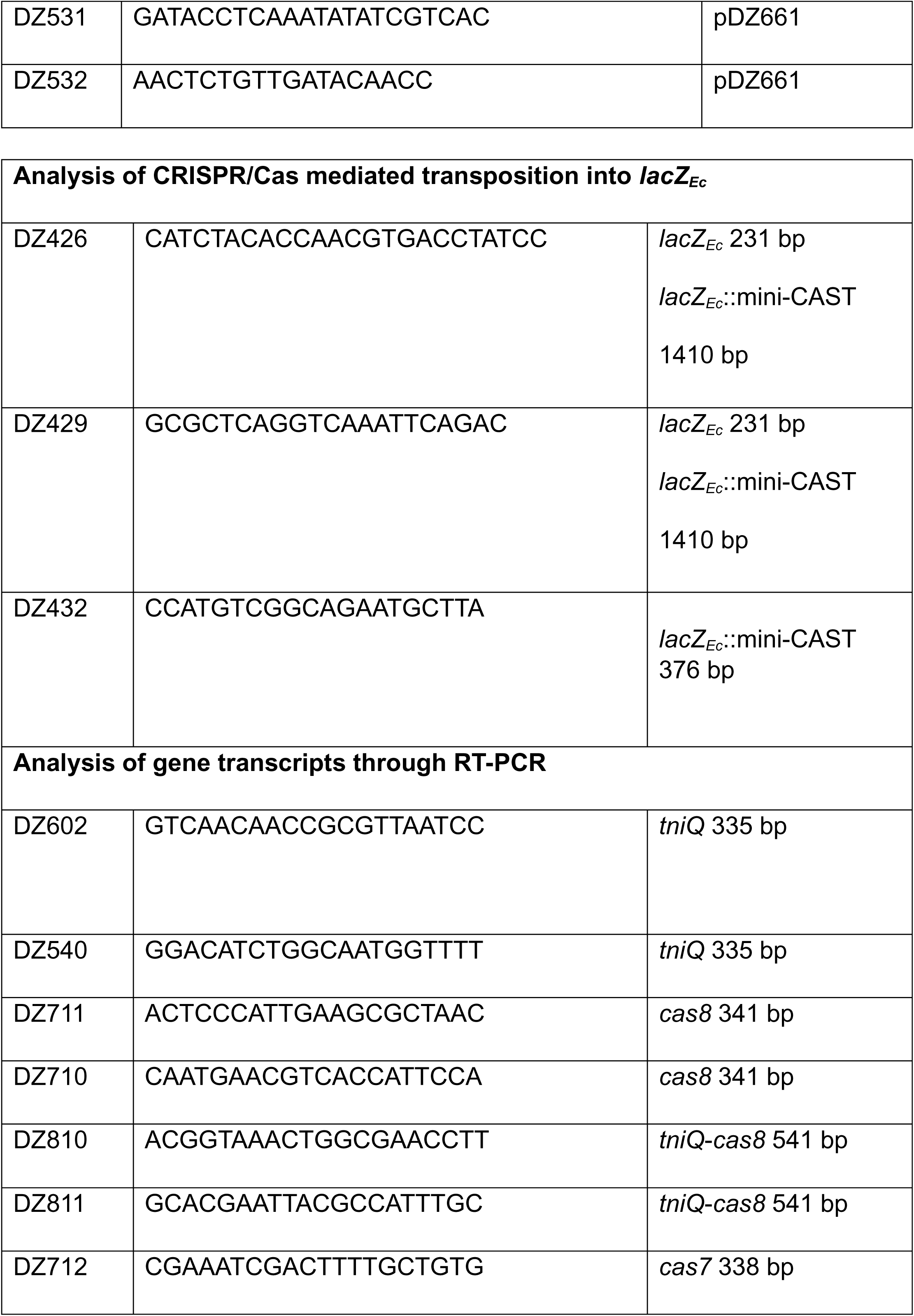
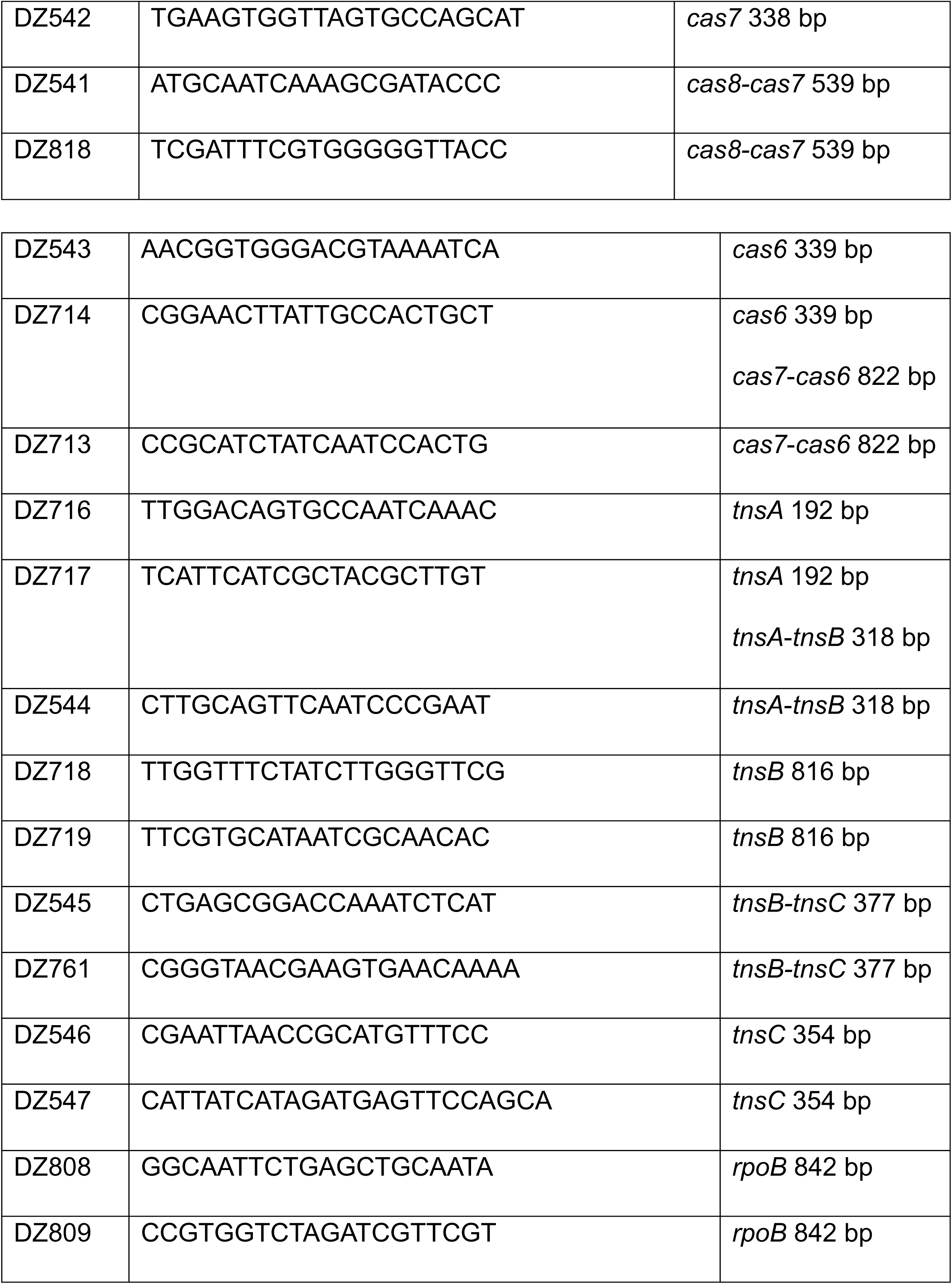
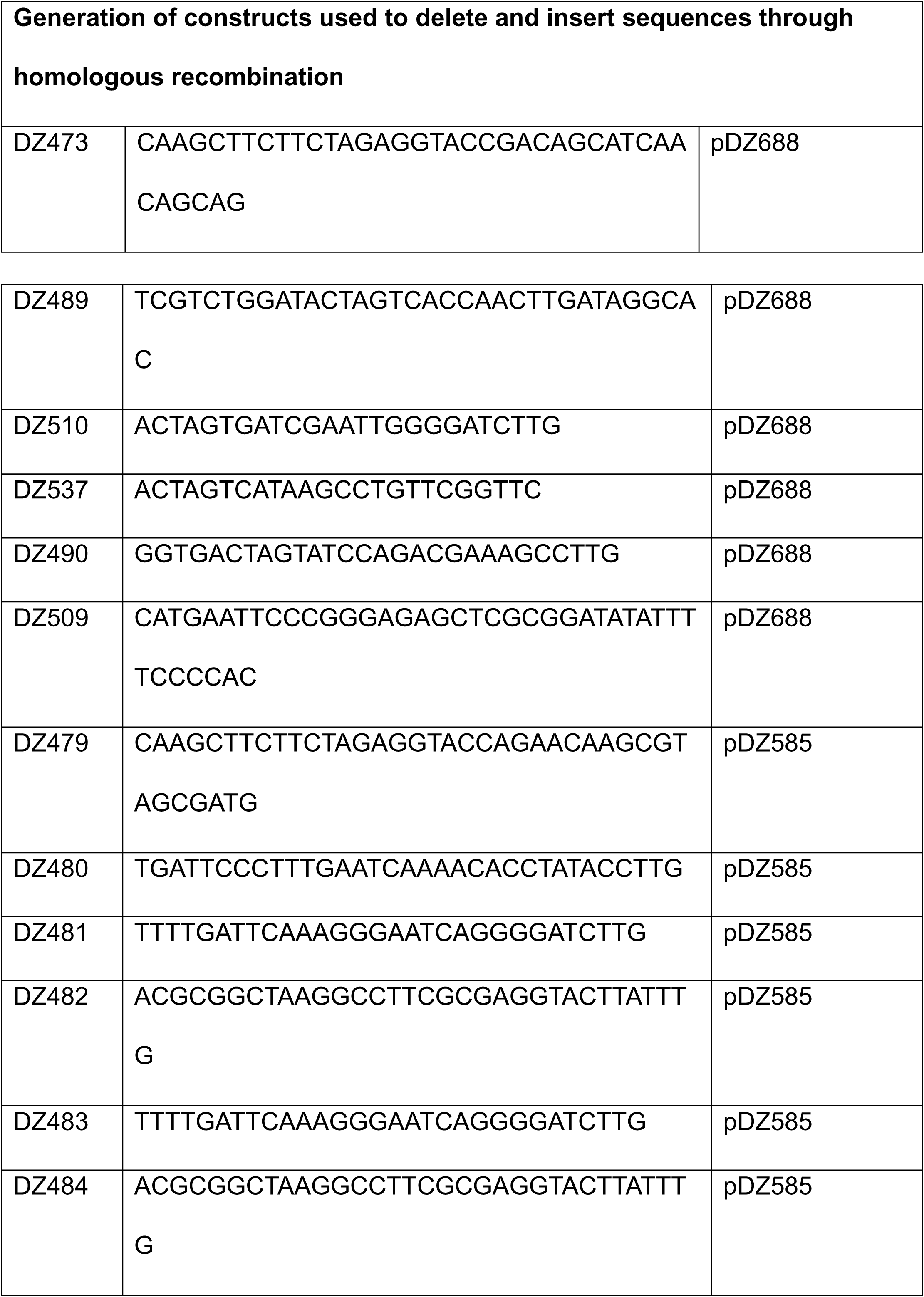
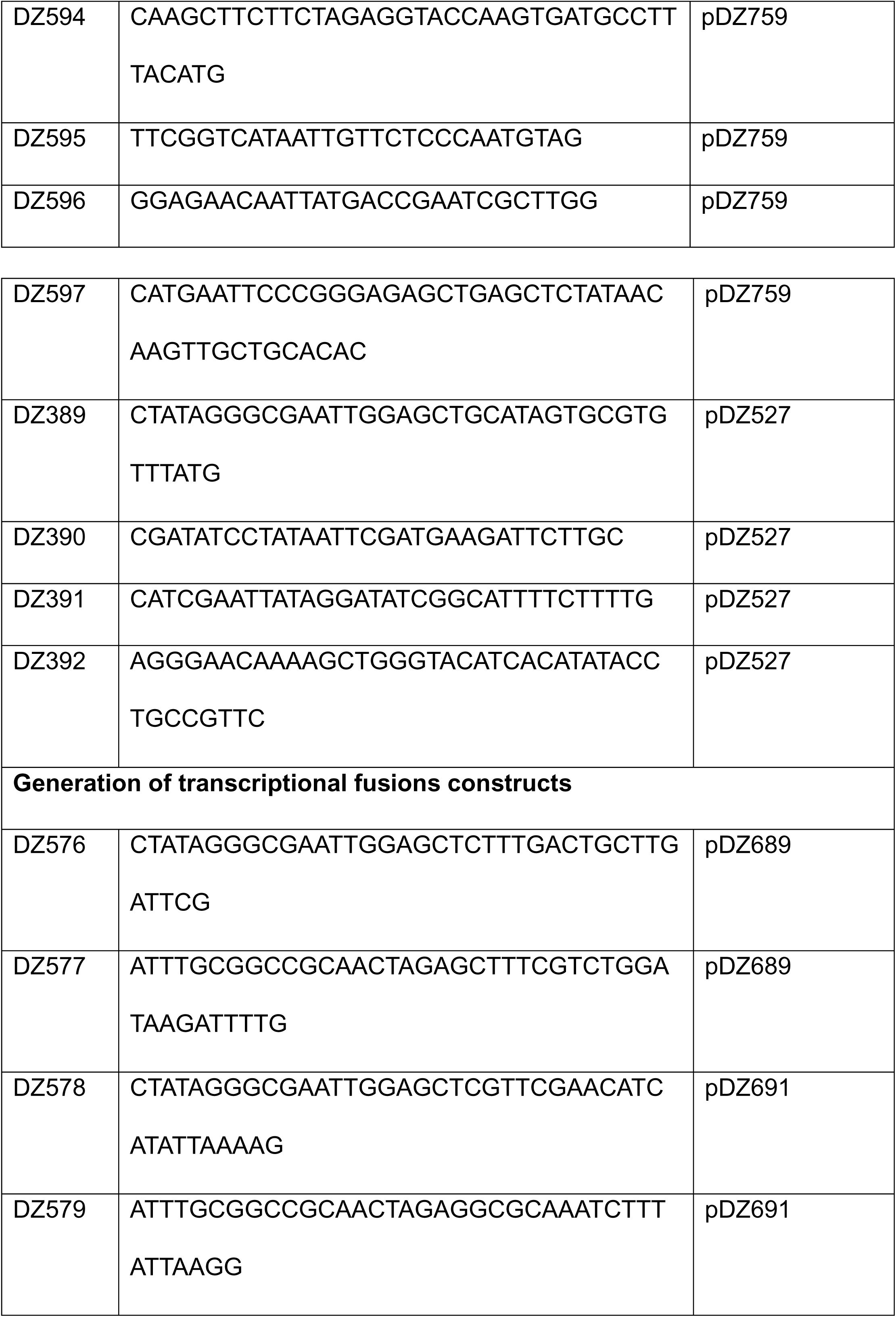
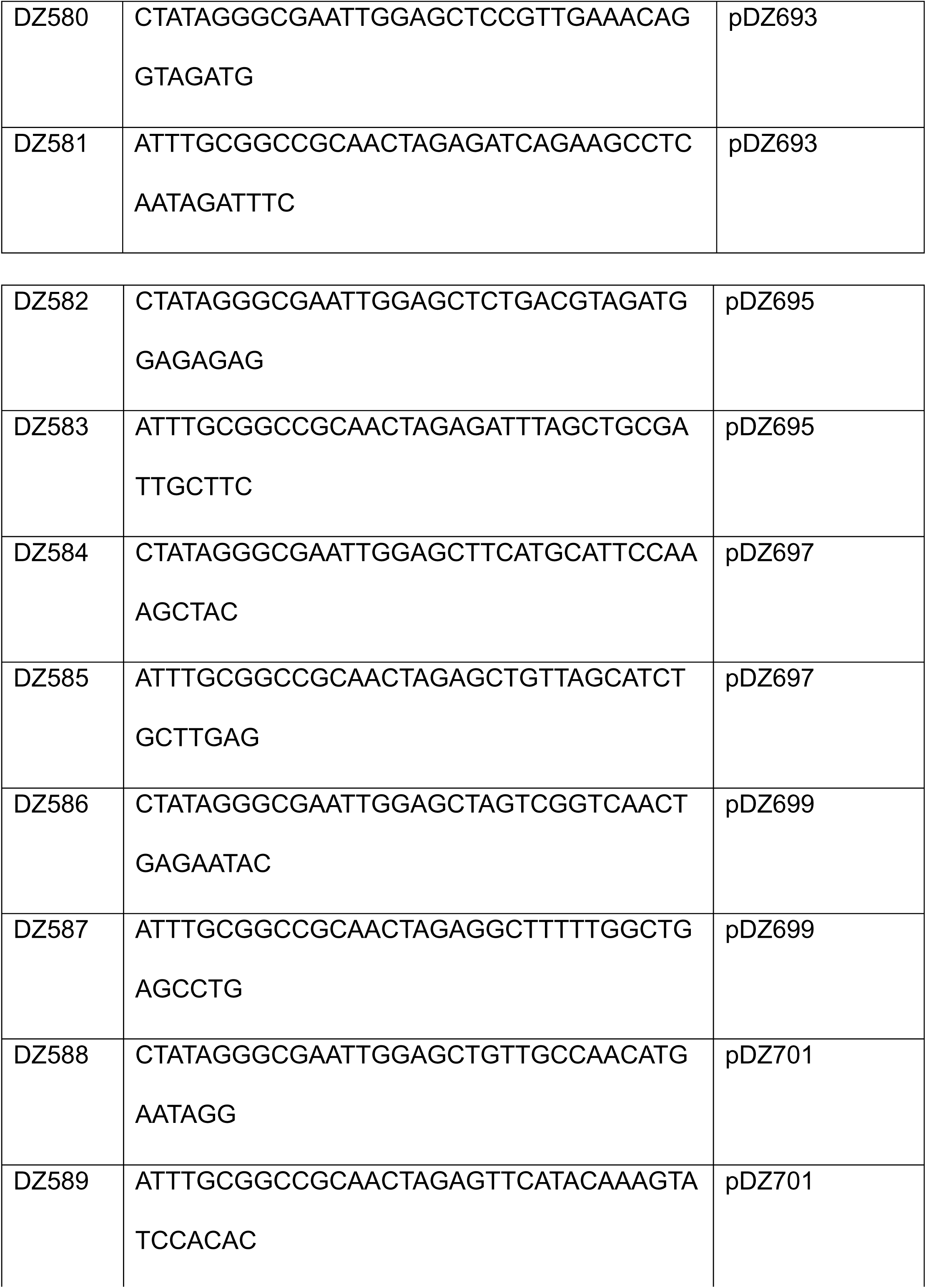
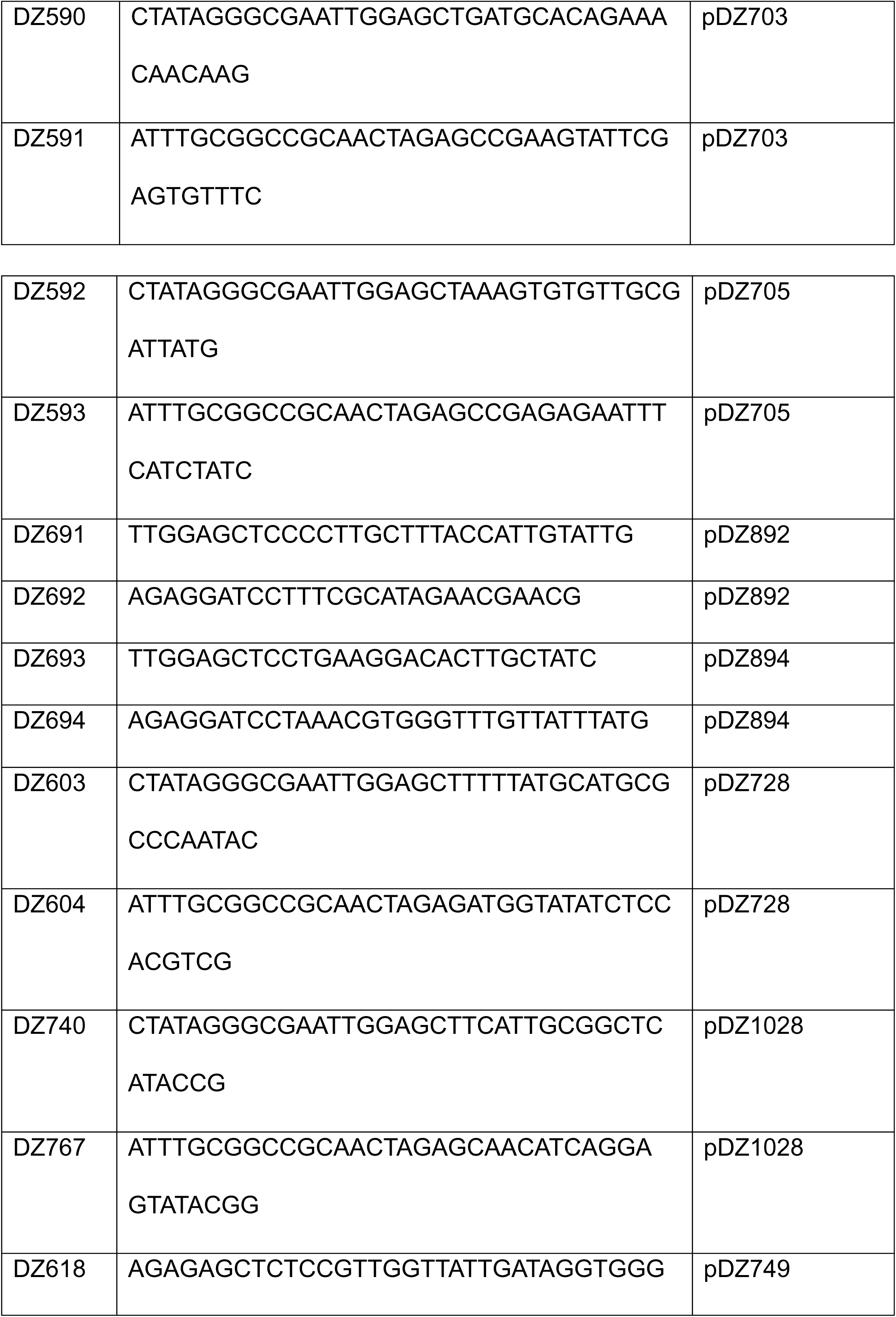
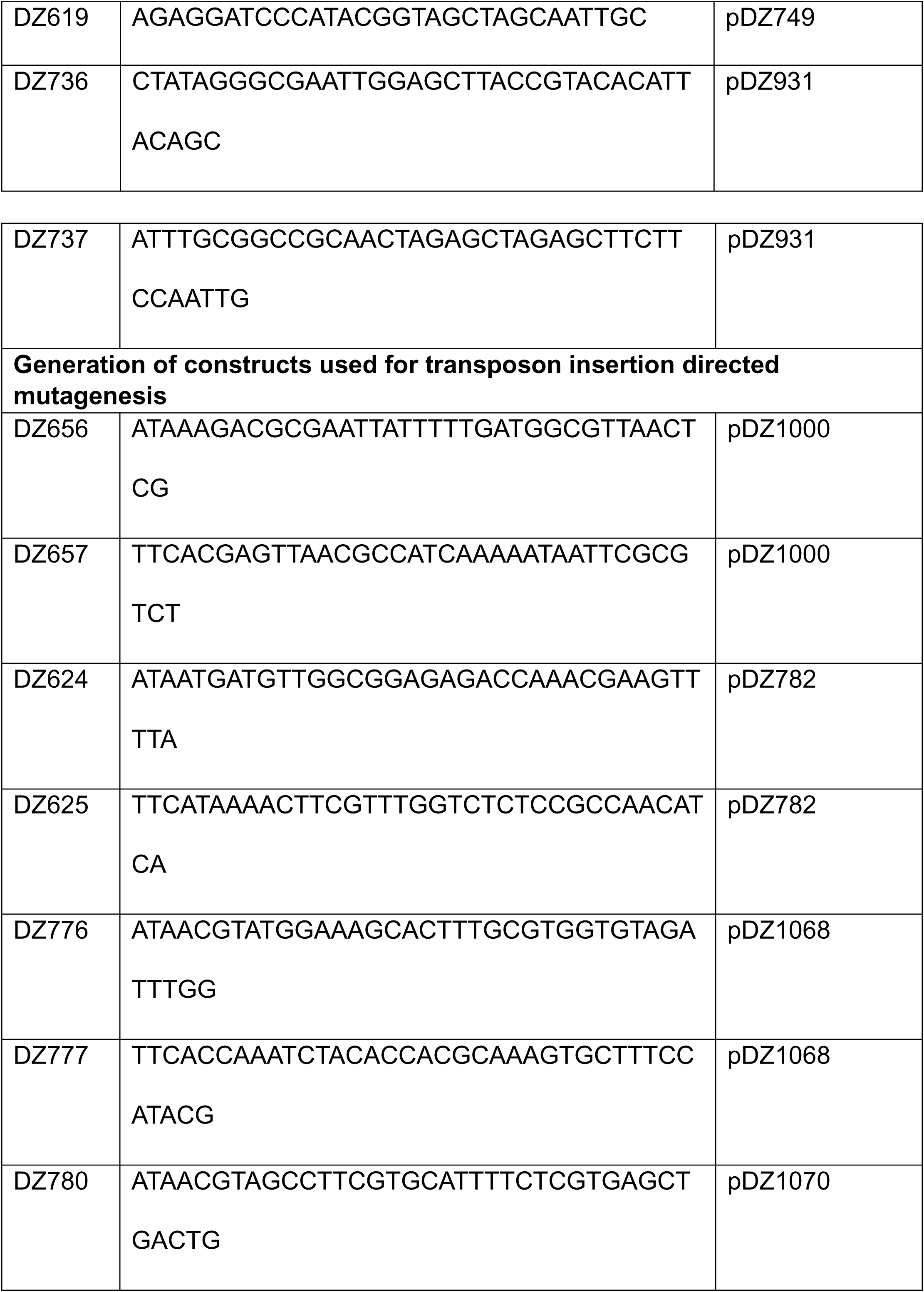
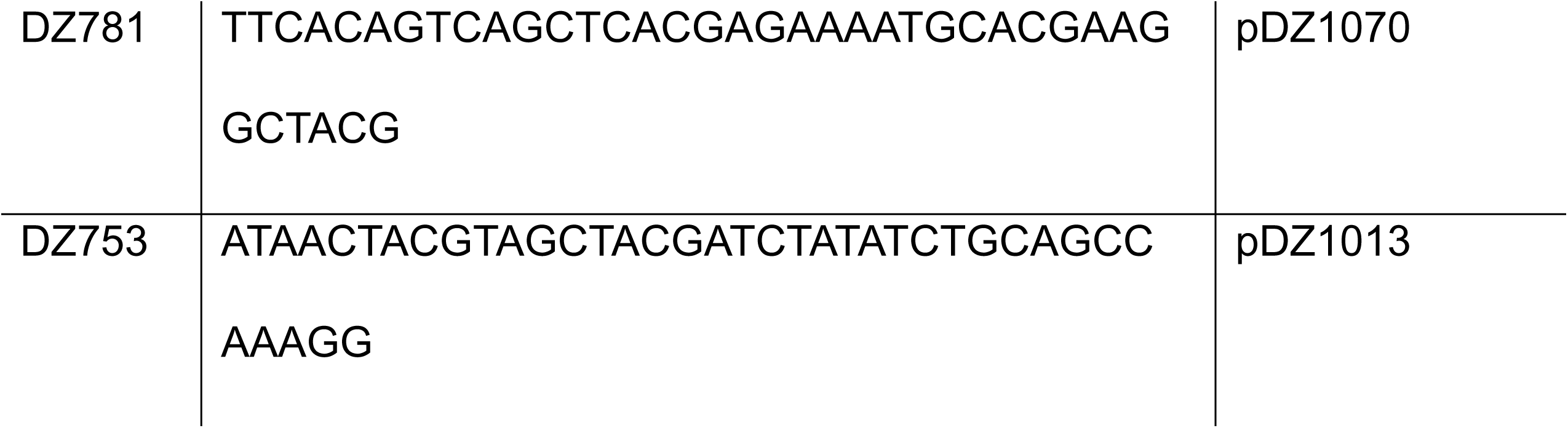
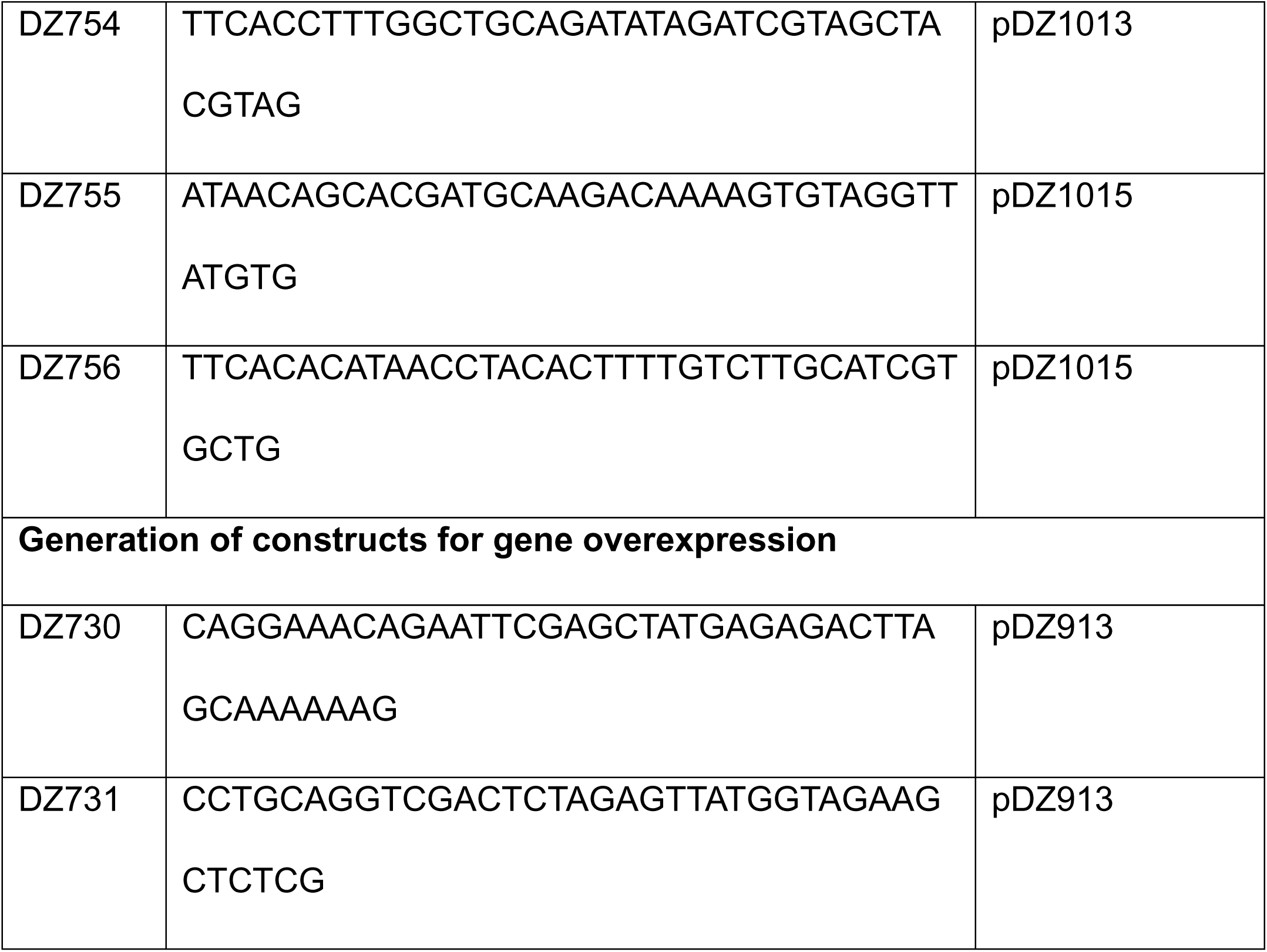
Primers used in this study.

The transcriptional fusions used in this study were generated by cloning upstream sequences from the genes of interest into the pBBRlux plasmid linearized with the restriction enzymes SacI-HF and BamHI-HF (New England Biolabs). Cloning was done either by isothermal assembly using the NEBuilder HiFi DNA Assembly Master Mix or by ligation with a T4 DNA ligase.

The construct for the overexpression of *vpxR* was generated in the backbone of the plasmid pMMB67EH-Gen. The *vpxR* coding sequence was cloned downstream of the Ptac promoter from pMMB67EH-Gen through isothermal assembly.

The plasmid containing the *Vpa*INTEGRATE system (pDZ879) was generated from intermediary constructs and assembled sequentially in a modular fashion. The plasmid pSL1142 (*Vch*INTEGRATE) Addgene #160730, a gift from Samuel H. Sternberg, was used as the backbone to assemble the components of the *Vpa*INTEGRATE system. We first PCR amplified the CRISPR array, the *tnsABC* loci and the cargo from *Vch*INTEGRATE, to generate three modular constructs (pDZ535, pDZ583 and pDZ551) in the cloning plasmid pJET 1.2 Blunt. These constructs have restriction sites that can be used for the final assembly into the backbone of pDZ387 (pSL1142).

We used site-directed mutagenesis with the Q5 site-directed mutagenesis kit from New England Biolabs to convert the *Vch*CAST typical CRISPR array in plasmid pDZ535 into the *Vpa*CAST typical CRISPR array generating plasmid pDZ633. To generate plasmid pDZ653 we cloned the *tniQcas876* genes from the *Vpa*CAST system in pDZ633 by means of an isothermal assembly reaction. The reaction consisted of the NEBuilder HiFi DNA Assembly Master Mix (New England Biolabs) and two PCR fragments, one corresponding to *tniQcas876* and the other corresponding to linearized pDZ633 obtained by inverted PCR. We added an ApaI restriction site downstream of *cas6* in pDZ653 to facilitate modular cloning of the *Vpa*CAST-*tniQcas876* region. Since the *Vpa*CAST *tniQ* gene has a an internal BsaI restriction site that would interfere with the cloning of CRISPR spacers, we removed this restriction site by site directed mutagenesis. We introduced a silent mutation in the codon for serine 231 in *Vpa tniQ* changing it from tct to tca.

Plasmid pDZ744 was generated by exchanging the *Vch tnsABC* genes with the *Vpa tnsABC* genes into the linearized pDZ583 plasmid. We used a T4 polynucleotide kinase from New England Biolabs to phosphorylate the 5’ ends of the PCR products corresponding to *Vpa tnsABC* and linearized pDZ853. The phosphorylated PCR products were ligated using a T4 ligase from New England Biolabs. The *Vpa tnsABC* fragment in pDZ744 have NdeI and ApaI restriction sites upstream of *tnsA* that allow the cloning of the *Vpa tniQcas876* module through the ligation of sticky ends.

Plasmid pDZ661 was generated by replacing the *Vch*CAST R and L-terminal sequences from plasmid pDZ551 with the *Vpa*CAST R and L-terminal sequences, sequentially, using a similar blunt-cloning approach as the one described for the generation of pDZ744.

To assemble plasmid pDZ879 containing the *Vpa*INTEGRATE system we first cloned the *Vpa* mini-CAST cargo into pDZ387 using Bsu36I and KpnI restriction sites to generate plasmid pDZ681. We next cloned the *Vpa tnsABC* module using NdeI and Bsu36I restriction sites within pDZ681 to generate pDZ799. Finally, pDZ799 was digested with the restriction enzymes SalI and ApaI to ligate the *Vpa* CRISPR-*tniQcas* fragment.

The allelic exchange of *lacZ_Vpa_* for *lacZ_Ec_* was done through double homologous recombination. First the *lacZ_Ec_* allele was PCR amplified using genomic DNA of BL21 DE3 and cloned into a linearized pMMB67EH-Gen plasmid (pDZ510). This construct was used as template to amplify the P_tac_-*lacZ_Ec_* region that was used for allele replacement. The suicide plasmid pDZ260 was used for allele replacement. This plasmid has 500 bp sequences that correspond to upstream and downstream homology regions flanking the *lacZ_Vpa_* allele. In between this homology regions are an FRT-*kanR*-FRT cassette followed by unique SacI and SpeI restriction sites. The Ptac-*lacZ_Ec_* region was cloned by isothermal assembly into pDZ260 linearized with SacI and SpeI to generate pDZ518. Allele exchange was achieved following the same protocol previously reported for the generation of gene deletions by double homologous recombination in *V. parahaemolyticus* (53, 54). The Kanamycin-resistance cassette was removed by site-specific recombination using plasmid pDZ514 that expresses the Flp recombinase and the counter selection allele *sacB*. To make pDZ514 and pDZ527 the *flp* cassette was amplified from plasmid pCP20 (55) and the *sacB* cassette from pRE118. The *flp* cassette consists of the *flp* gene and a gene that encodes a temperature-sensitive λ repressor gene. Both genes are controlled by a single λ promoter. The *flp* cassette and the *sacB*cassette were assembled into either a pBBR1MCS-5 (pDZ514) or pBBR1MCS-2 (pDZ527) backbone through isothermal assembly. Site specific recombination was induced by incubating exponentially grown cells at 42 °C for 1 to 2 hours. The desired eliminations were validated by PCR.

The deletion of the *Vpa*CAST system from chromosome 2 in *V. parahaemolyticus* RIMD2210633 was achieved by a combination of homologous and site-specific recombination approaches. The L-terminal sequence was eliminated by double homologous recombination using plasmid pDZ759. To generate pDZ759 regions of approximately 500 bp flanking the L-terminal sequence from VpaI-7 were cloned into the suicide plasmid pRE118. Deletion of the L-terminal sequence from VpaI-7 by double homologous recombination was achieved as previously reported (53, 54). To delete a region from VPA1387 (*tniQ*) to VPA1397 (*yciA*) an FRT-*genR* gentamicin resistance cassette was inserted upstream of *tniQ* and an FRT-*sfgfp* cassette was inserted within VPA1397. This was achieved by double homologous recombination following a similar approach as the one used to delete the L-terminal sequence. Flp site-specific recombination was used to delete the region flanked by the FRT-*genR* and the FRT-*sfgfp* cassettes.

### Luminescence assay

The luminescence assays in planktonic cultures were performed as previously described (53, 54). To quantify light production in planktonic cells, we diluted cultures grown overnight 1:200 in 5 mL of LB broth in test tubes with a volumetric capacity of 20 mL and incubated them under shaking conditions (200 rpm) at 30°C until reaching an optical density at 600 nm (OD600) between 0.6 and 0.8. Light production from cells transferred to white 96-well plates was detected using a Synergy H1 plate reader from BioTek. The relative luminescent units (RLU) are calculated as arbitrary luminescence per mL and divided by the OD600 of the cultures.

### Generation of a consensus logo for the putative binding site of VpxR

To find conserved motifs we used 300 nucleotides of the upstream sequence of *vpxR* and orthologues from *Aliivibrio salmonicida* LFI1238*, Vibrio vulnificus* CMCP6, *Vibrio campbellii* ATCC BAA-1116, and *Vibrio cholerae* O395 and 84 nucleotides of the upstream sequence of the *Vpa*CAST atypical CRISPR array and homologous atypical CRISPR arrays identified in the genomes of *Vibrio diabolicus* FDAARGOS 105 and *Vibrio natriegens* CCUG 16374. Motifs and their consensus logo were generated using the Multiple Em for Motif Elicitation (MEME) algorithm (56).

### Generation of reprogrammed CRISPR arrays in the INTEGRATE systems

Plasmids pDZ387 and pDZ879 have a non-targeting spacer sequence with two internal BsaI restriction sites that can be used to clone targeting spacer sequences. The targeting spacer sequences were designed with a python script generated by the Sternberg Lab and available through Github (https://github.com/sternberglab/CAST-guide-RNA-tool). The targeting spacers are 32 bp long and where cloned as oligoduplexes in the backbone of pDZ387 or pDZ879. The oligoduplexes were generated by annealing complementary oligonucleotides. The annealing procedure was done in a 100 μL solution with each oligonucleotide at a 20 μM concetration in a buffer composed of 60 mM KCl, 6 mM HEPES-pH 7.5, and 0.2 mM MgCl2. The solution was incubated a 95°C for 1 minute and then cooled down to 25°C by reducing the temperature 1°C every minute in a Thermocycler. The annealed oligoduplexes were phosphorylated using a T4 polynucleotide kinase from New England Biolabs. The plasmids pDZ387 and pDZ879 were digested with BsaI-HFv2 from New Engalnd Biolabs. The phosphorylated oligoduplexes were ligated to the linearized pDZ387 and pDZ879 plasmid using a T4 DNA ligase.

### INTEGRATE mediated transposition assays

To use a conjugation approach to mobilize the pDZ879, pSL1142, pDZ952 and pDZ1000 plasmids to both *E. coli* and *V. parahaemolyticus* strains, we generated an *E. coli* BL21 DE3 receiver strain with a selection marker. To do so, we introduced into *E. coli* BL21 DE3 the plasmid pBAMD1-6 (57) which has a mini-Tn5 transposon with a gentamicin resistance cassette. We selected a gentamicin resistant colony (EcDZ1067) with a random transposition event that grew like the parental strain. The pSL1142 derivative plasmids were horizontally transferred to the EcDZ1067 and VpDZ838 strains by a triparental mating approach using the helper plasmid pRK600 as previously reported (54). EcDZ1067 derivative transconjugants were selected in LB-agar plates with gentamicin and kanamycin while VpDZ838 derivative transconjugants were selected in LB-agar plates with streptomycin and kanamycin. *E. coli* and *V. parahaemolyticus* strains were incubated at 30°C for 48 and 24 hours, respectively. X-gal at 40 μg/mL and IPTG at 0.1 mM were added to LB-agar plates when indicated. The blue vs white colony count was done from in average 150 colonies per plate.

Insertion mutations using the *Vch*INTEGRATE system from plasmid pSL1142 (pDZ387) were generated similarly as described above. The pDZ387-derivative plasmids with a reprogrammed targeting spacer were mobilized into *V. parahaemolyticus* RIMD2210633 through triparental mating. Transconjugants were selected in LB-agar plates with streptomycin and kanamycin and incubated for 48 hours at 30°C. The mini-CAST insertion was analyzed with specific primers. The pDZ387-derivative plasmids were curated by growing the insertion mutants without selection twice. Kanamycin sensitive colonies were genotyped by colony PCR using a MyTaq polymerase Mix from Meridian Bioscience.

## Acknowledgements

This work was financially supported by DGAPA-PAPIIT (UNAM) project no. IA203323 awarded to David Zamorano-Sánchez. Jessica Cruz-López received a scholarship financed with DGAPA-PAPIIT (UNAM) project no. IA203323. Jesús E. Alejandre-Sixtos is conducting his doctoral studies in the “Programa de Doctorado en Ciencias Biomédicas”, Universidad Nacional Autónoma de México (UNAM) and has received CONAHCYT fellowship 1242669.

Eugenio Lopez-Bustos, Paul Gaytan, Santiago Becerra, and Jorge A. Yañez (Unidad de Síntesis y Secuenciación, Instituto de Biotecnología-UNAM) provided technical assistance with oligonucleotide synthesis and DNA sequencing.

